# Consensus prediction of cell type labels with popV

**DOI:** 10.1101/2023.08.18.553912

**Authors:** Can Ergen, Galen Xing, Chenling Xu, Michael Jayasuriya, Erin McGeever, Angela Oliveira Pisco, Aaron Streets, Nir Yosef

## Abstract

Cell-type classification is a crucial step in single-cell analysis. To facilitate this, several methods have been proposed for the task of transferring a cell-type label from an annotated reference atlas to unannotated query data sets. Existing methods for transferring cell-type labels lack proper uncertainty estimation for the resulting annotations, limiting interpretability and usefulness. To address this, we propose popular Vote (popV, https://github.com/YosefLab/popV), an ensemble of prediction models with an ontology-based voting scheme. PopV achieves accurate cell-type labeling and provides effective uncertainty scores. In multiple case studies, popV confidently annotates the majority of cells while highlighting cell populations that are challenging to annotate. This additional step helps to reduce the load of manual inspection, which is often a necessary component of the annotation process, and enables one to focus on the most problematic parts of the annotation, streamlining the overall annotation process.

## 1 Introduction

Cell type annotation is a crucial task in analyzing single-cell RNA sequencing data. The quality of the annotations has a direct impact on downstream analyses such as the comparison of cell type composition, as well as the analysis performed on a per cell type basis [1]. Manual annotation is highly time-consuming and requires tissue-specific and sequencing technology-specific domain knowledge. Thus, as scRNAseq becomes an increasingly standard lab technique, there is a growing need to generate automated annotations. We propose here the use of a collection of cell type prediction models to provide not only automated annotations but also well-calibrated measures of uncertainty enabling effective incorporation of the human-in-the-loop component of the annotation process.

Automated cell type annotations encounter several challenges [2]. There is no gold standard ground truth for cell type annotation within a specific data set. Biology is complex, and when cell states vary continuously, delineations between cell types are imprecise, and even human experts may disagree on the exact phenotype of a specific cell. Therefore, it is essential that annotation methods highlight areas of uncertainty that require expert knowledge input. The continuous nature of cell states [3], along with stochasticity in the sequencing process, as well as the domain knowledge of the person manually annotating the data set, can lead to cells being annotated at varying levels of specificity even within the same data set. Across multiple data sets, factors, like continuous discovery and identification of novel cell subtypes, lead to discrepancies in cell type identification. There are a plethora of automated cell type annotation methods [4]. However, differences in cell type granularity, experiment-specific nuisance factors, and technology-dependent sparsity of gene expression lead to no clear ‘best method’ for automatic annotation. Based on these factors, we pose that it is crucial for automatic cell type annotation pipelines [5] to: highlight areas of uncertainty that may require manual scrutiny; balance specificity of predictions with accuracy; be easily accessible and usable.

To address these challenges, we developed popularVote (popV), a flexible and scalable automated cell type annotation framework that takes in an unannotated query data set from a scRNAseq experiment, transfers labels from an annotated reference data set, and generates predictions with a predictability score indicating the confidence of the prediction. We pose here that various prediction methods will disagree in their prediction if an annotation is not accurate, whereas they will tend to agree if the predicted cell type is the correct one. We named our method popularVote because instead of relying on the predictions of a single classifier, popV takes a consensus approach and incorporates the predictions from eight automated annotation methods. PopV also takes into account annotations at different levels of granularity by aggregating results over the Cell Ontology [6]; an expert-curated formalization of cell types in a hierarchical structure with a standardized vocabulary. PopV is available as an easy-to-install, open-source Python package and is designed to be a flexible framework for incorporating future cell type classification methods. We provide a notebook that allows the prediction of new data sets and provide pre-trained models for 20 different organs based on The Tabula Sapiens data set [7].

## 2 Results

### Overview of popV

PopV takes a consensus of experts approach to the task of automated cell type annotation. The input is an unannotated query data set together with an annotated reference data set (Figure 1A). Both data sets are expected to contain raw count data and demonstrate that popV can be applied to UMI, as well as non-UMI-based technologies. PopV then runs eight different annotation methods: random forest (RF) [8], support vector machine (SVM) [8], scANVI [9], OnClass [10], Celltypist [11] and *k* nearest neighbors (*k* NN) after batch correction with three single cell harmonization methods: scVI [12], BBKNN [13], and Scanorama [14] (Figure 1). The eight prediction algorithms were chosen because they were shown to have good prediction accuracy [15] and/or good harmonization performances [16]. These methods encompass supervised methods that are trained only on labeled data (Random Forest - RF, Support Vector Machine - SVM, OnClass, Celltypist, *k*-nearest neighbor classifier (*k*NN) after applying unsupervised harmonization methods that are agnostic to label information during training (BBKNN, Scanorama, scVI) and a semi-supervised method trained with both labeled and unlabeled data (scANVI). The unsupervised methods are coupled with a *k*NN classifier to generate label predictions. However, we emphasize that popV offers an intuitive application interface (API) for rapid inclusion of additional annotation methods. We demonstrate this capability through a code snippet for adding a new classifier (*k*NN after batch correction with harmony [17]) in the Methods section.

**Figure 1:**
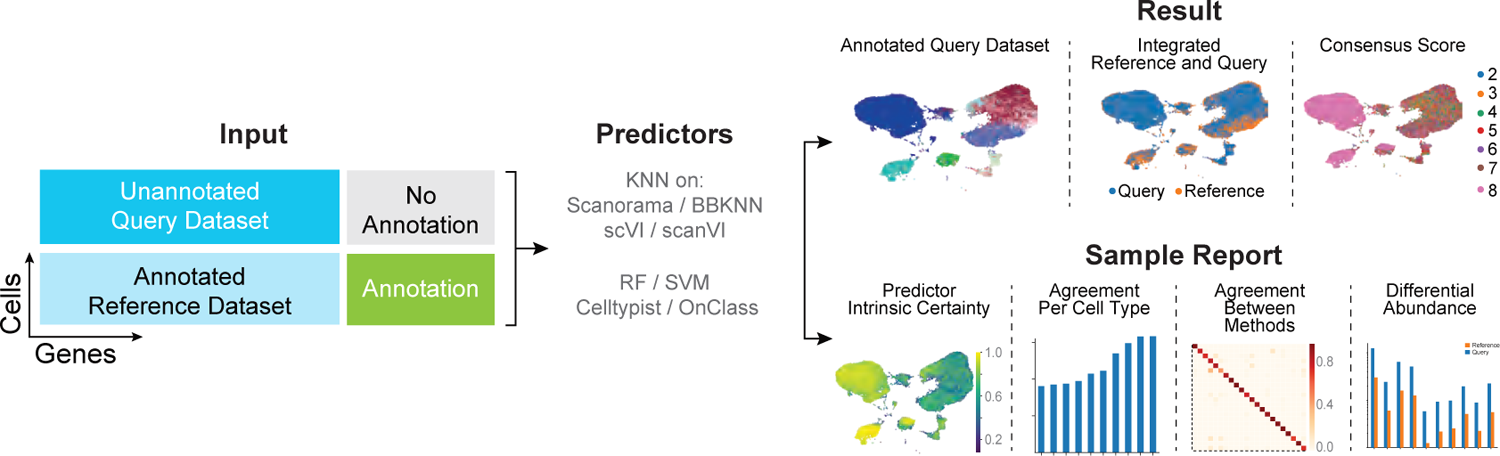
Framework of popV for automatic cell type annotation. PopV takes as input an unannotated query data set and an annotated reference data set. Each expert algorithm predicts the label on the query data set to yield a cell type annotation. The certainty of the respective label transfer can be quantified by scoring the agreement of those methods. The workflow yields a sample report to provide the user with insights into the annotated labels.

After applying each of these methods separately, popV proceeds to aggregate the resulting predictions for two purposes (Suppl. Figure 1). The first is to designate a single ‘consensus’ annotation for every query cell. The second purpose is to quantify our certainty in this prediction. We estimate the consensus annotation using a simple majority vote procedure, counting for each annotation label the number of algorithms that support it. The one exception to this simple procedure is how we treat OnClass annotations, as OnClass is the only method in this collection that is capable of predicting cell types that do not exist in the reference data set. It does so through a two-step process - first selecting an annotation out of the collection of labels in the reference data set and then propagating it to identify a potentially more precise label in the Cell Ontology (even if this label is absent from the reference). To account for these “out of sample” cell type annotations, we consider every label that is on the path from the root of the ontology down to the OnClass-predicted label as a predicted label (Suppl. Figure 1). We then perform majority voting across all labels (with OnClass having multiple “votes” at different levels of hierarchy). We have attempted using a simple majority vote with the “within sample” annotation from the first stage of OnClass, and with no propagation along the Cell Ontology. In most of our analyses, we found our first strategy to be preferable. Throughout the manuscript, we report this method, except that it is highlighted differently.

A potentially useful property of many of the algorithms included in popV is an “algorithm-intrinsic” estimation of prediction certainty. This could, in principle, be leveraged to compute a weighted consensus. However, we found that the certainties are calibrated differently for the different methods, which makes this approach futile as it will weigh more on the predictions of the most presumptuous classifiers.

After calculating the consensus score, popV generates a sample report that includes prediction summaries as well as integrated views of the query and reference data sets (see Suppl. Report 1). For the latter, it displays UMAPs for the joint visualization of the reference and query data sets for the four methods that perform data integration (Figure 1), as well as a bar plot comparing cell type frequencies in the reference and query data set to highlight the differential abundance of various cell types. One set of summaries in the report are confusion matrices between the consensus predictions and each individual method to indicate which cell types were confused with another cell type for any particular method. The report also includes a per cell type display of the consensus score (i.e., number of agreeing methods - between 1 and 8) to highlight which cell types are overall difficult to predict. Complementing this ‘algorithm-extrinsic’ estimation of certainty, we also output the intrinsic uncertainty (i.e., classifier score) of each of the eight methods (these scores are defined in the method section). We emphasize that intrinsic and extrinsic uncertainty are two complementary measurements essential to quantify the performance of a set of cell annotation tools.

To allow for fast annotation of new query data sets, we provide pre-trained models for all 20 organs present in Tabula Sapiens [18]. Pre-training is possible for all methods except Scanorama and BBKNN, which compute a joint embedding of reference and query data set and make pre-training infeasible. For scVI and scANVI, we provide pre-trained embeddings of the reference data set and map the query data set to this embedding using scArches [19]. PopV has three different modes of cell type prediction; in *retrain* mode all classifiers are trained from scratch, which requires an hour for 100k cells in a Google Colab session, in *inference* mode pre-trained models are used where applicable, which requires 30 minutes for 100k cells, in *fast* mode only pretrained models are used and only cell types in the query data set are predicted, which requires 5 minutes for 100k query cells. PopV is available as an open-source Python project and includes an online Google Colab notebook with free computing resources. The codebase enables the addition of new reference data sets (in addition to Tabula Sapiens) through a simple API and can be invoked from the same notebook environment. We recommend that in any newly added reference data sets, the annotations should be consistent with Cell Ontology, either by matching terms in the ontology or by hierarchically assigning new terms to existing terms in the ontology. To this end, we provide scripts to add custom cell type labels to The Cell Ontology before processing by popV (where it is used for running OnClass and calculating our consensus scores).

### PopV prediction score discriminates high and low quality annotations

We evaluated the performance of cell type annotation using popV with a human lung cell atlas as the query data set [20] and the lung tissue of Tabula Sapiens as a reference data set. The Lung Cell Atlas is carefully annotated to a high level of granularity. It contains a wide variety of cell types across immune cells, epithelial cells, endothelial cells, and stromal cells and is therefore well suited for studying tissues with diverse labels. To make the labels comparable across both data sets, we translated the Lung Cell Atlas labels to the corresponding terms in the Cell Ontology (Suppl. Figure 2).

PopV achieves high accuracy on the Lung Cell Atlas. We visualize the popV predictions against the manual annotations in the Lung Cell Atlas and see a strong agreement between the prediction and the original annotation, as well as a good integration between the query and the reference cells (Figure 2A). We decided here to use scANVI integration as it showed the highest performance in scIB metrics, which measures data integration and biological conservation [16] (Suppl. Figure 3A). To evaluate the quality of our predictions, we compute accuracy terms based on the Cell Ontology tree (see Methods). *Exact Match*, as the name implies, means that the predicted cell type is exactly the same as the manual annotation. Furthermore, intuitively, a prediction algorithm that predicts one cell type as another similar cell type performs better than a prediction algorithm that predicts the cell is of an unrelated type. The *Parent Match*, *Child Match* and *Sibling Match* take this into account and measure if the predicted cell type is the parent, child, or sibling in the Cell Ontology tree compared to the ground truth annotation. This measure is especially useful if a cell type label exists only in the query and not in the reference data set. Every prediction that did not match any of these relationships was classified as *No Match*. PopV overall achieves high accuracy for most cell types (Figure 2B and Suppl. Figure 3C). Except scANVI and OnClass, all methods have comparable performance in this data set. Furthermore, we compared the performance of popV with Seurat, which is another popular tool for cell type annotation transfer [21] and found that Seurat performs worse than most methods used in popV. We also included OnClass predictions after step one (OnClass_seen), where OnClass only predicts cell types that were present in the reference data set and found this to perform similarly to the good-performing annotation tools, so that the lower performance of OnClass here is solely due to the prediction of unseen cell types. Overall, popV performed best for the number of *Exact Match* and comparable in the number of cells with *No Match*. Overall, the popV prediction is more accurate than any of the single methods. For a better insight into the prediction, we display bar plots in the report for popV highlighting the abundance of cell types in query and reference data set, as well as prediction accuracy (Suppl. Figure 3) and display confusion of cell types using alluvial plots (Suppl. Figure 4). We used this case study to demonstrate the utility of our consensus scoring strategy, compared to a simple majority vote, observing an overall superior performance (Suppl. Figure 5).

**Figure 2:**
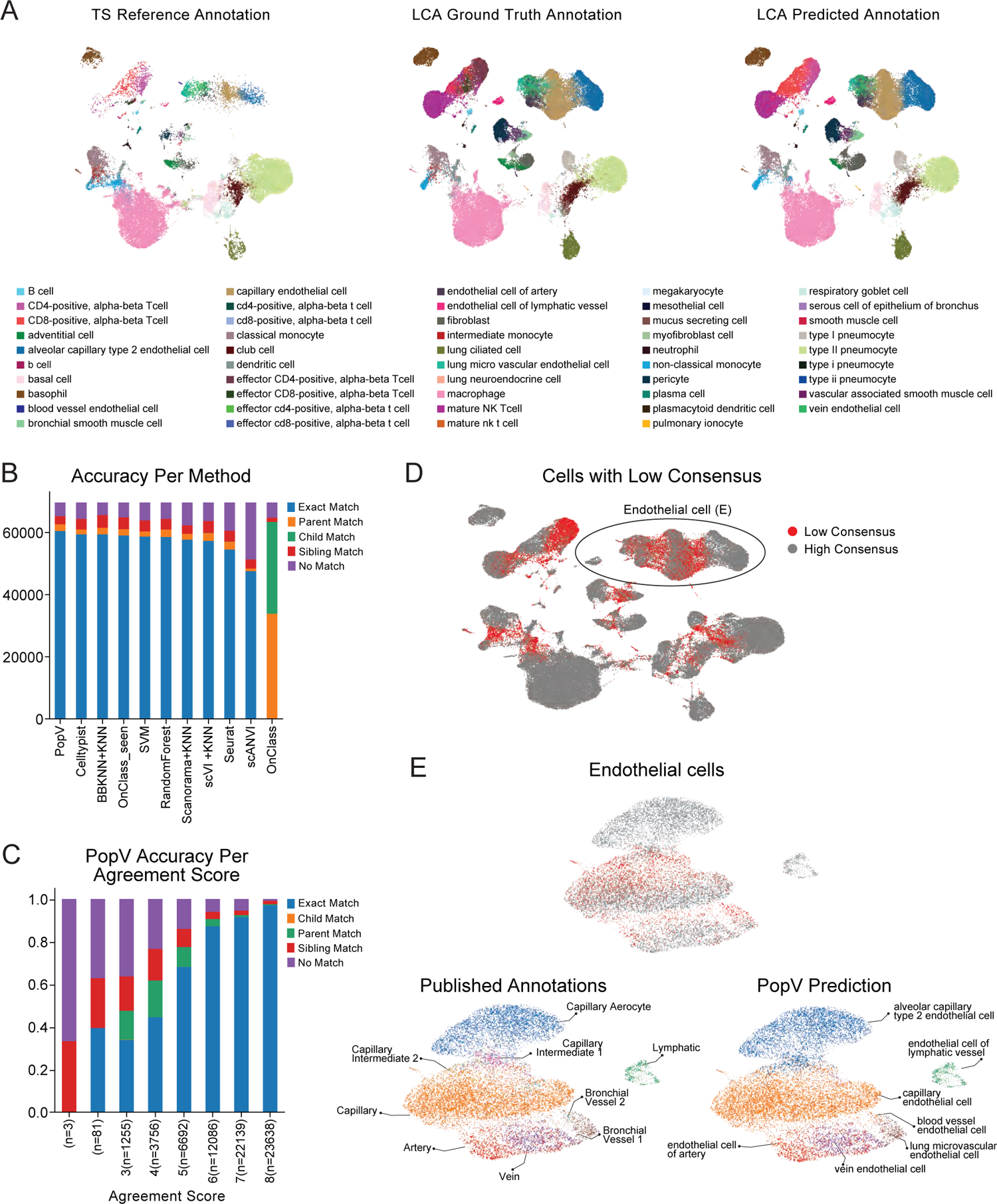
PopV prediction on Lung Cell Atlas and Tabula Sapiens lung as reference is accurate and interpretable. **A**, UMAP embedding after scANVI integration of Tabula Sapiens reference cells, Lung Cell Atlas query cells labeled with predicted label and Lung Cell Atlas query cells labels with the ground-truth label. **B**, Ontology accuracy (see methods) for the various methods computed on the query cells. **C**, Ontology accuracy for the prediction scores in popV. **D**, highlighted cells with a consensus score of 4 or less (Low Consensus) and (E) Zoomed in view of endothelial cells in the Lung Cell Atlas with popV predicted labels and ground-truth labels displayed. The zoomed-in picture is rotated by 90 degrees to allow readability of all labels. *Alveolar capillary type 2 endothelial cell* is the cell ontology term for *capillary aerocytes*. The Lung Cell Atlas annotated additional cell types between *Capillary Aerocytes* and *Capillary*

When checking the popV prediction scores, we found that the accuracy of the prediction is highly correlated with the prediction score (Figure 2C). For scores of 6 and higher, we found that more than 90% of the annotations were exact matches with the ground truth. For scores of 8, which is a perfect agreement between all methods, 98% of the predictions were exact matches. For scores of 3 and lower, the prediction accuracy was lower than 50%, highlighting that the popV consensus score is a valuable metric to reflect the classification accuracy and points to groups of cells that should be further (and manually) scrutinized.

When considering cells that were assigned with a low consensus score, we found three possible reasons that may explain the disagreement between the different methods (Figure 2D). The first is that the distinction between certain cell subsets with different labels is unclear. This often arises in cases of a continuum of cell states with no clear decision boundary in transcriptome space. In such cases, the boundaries determined by different algorithms may vary (because they depend on different objectives or techniques), leading to low consistency. It is, however, exactly those cases that merit closer (and often manual) inspection, and - if needed - assignment of multiple optional labels. As an example, we found several areas of low consensus score in the various lung endothelial cells (Figure 2E). Most endothelial cells with a low consensus score arise between *capillary endothelial cells* and *alveolar capillary type 2 endothelial cells*. In this region, the various algorithms disagree on the correct boundary, but all algorithms predict those cells with either of those labels. We found that *alveolar capillary type 2 endothelial cells* express EDNRB and HPGD, and *capillary endothelial cells* express FCN3 and IL7R. Cells between both cell types are double positive in both markers, while they do not show any specific marker gene. Therefore, we conclude that neither the term *capillary endothelial cell* nor the term *alveolar capillary type 2 endothelial cell* is adequate to describe these cells, but their phenotype is between both cell types. Thus, it is a region that requires manual scrutiny to determine the correct label of those cells. In fact, such scrutiny was applied in the original annotation of the Lung Cell Atlas, which was not provided to popV and which labeled these cells as *Capillary Intermediate 1* and *2*. Therefore, this example demonstrates that a low consensus score can help identify areas that require a refined label, possibly extending the vocabulary available in the reference atlas.

The second reason for a low consensus score in this case study occurs when the query data set contains subsets of cells that are absent from the reference atlas. As an example, while the Lung Cell Atlas (which we use as the query) includes a subset of endothelial cells that was originally labeled *bronchial vessel 2*, this subset (and its respective label) seems to be absent from our reference atlas. Indeed, when checking marker genes for these cells, their expression was high in PLVAP and low in the vein endothelial marker ACKR1 (Suppl. Figure 6), which can be interpreted as an intermediate stage between *capillary endothelial cells* (negative for both markers) and *lung microvascular endothelial cells* (positive for both markers). This combination of marker gene expression was not observed in Tabula sapiens and therefore marks a cell type not present in the reference data set.

The third possible reason we find for the low consensus is inaccuracy in the reference annotation. As an example, we found a subset of T cells with a low consensus score (Suppl. Figure 7). All cells of this group that came from the query data set were originally labeled (by the authors of the Lung Cell Atlas) as *effector CD4 positive alpha-beta T cells*, while cells originating from the Tabula Sapiens reference were labeled with a mixture of CD4 and CD8 T cells. Consequently, most algorithms in popV labeled this low-consensus group as a mix of CD4 and CD8 T cells with different decision boundaries. Manually following up this low scoring group, we checked the marker gene CD8A and found a clear cut-off that distinguishes the CD4+ (T helper) and CD8+ (cytotoxic) subsets, in a manner consistent with the (hidden) query annotation. Despite this clear delineation, we found that many CD8 negative cells are labeled in the reference atlas as CD8+ T cells. A low consensus score in this group of cells helped to identify wrongly annotated cells in the reference data set, and we highlight that manual scrutiny can clean up these wrong labels.

We have demonstrated here that the consensus score can highlight regions that require manual scrutiny and that reannotation in those regions can be performed using marker gene expression. This leads to novel delineation of cell types not discovered in the reference data set, detection of query-specific cell types, and correction of the cell type label of wrongly assigned reference cell type labels.

### PopV detects a-priori differences between query and reference tissues and enables label transfer at different levels of granularity

We next set out to study cell type annotation in cases where the reference and query data sets originate from similar, yet not identical tissue environments. This will allow us to test whether the low-consensus regions of popV align with differences between the tissues and therefore require further scrutiny. To this end, we decided to use a recent multidonor effort to study the anatomy of the human brain [22]. In this study, three million nuclei were sequenced in three postmortem donors and approximately 100 dissections in various regions of the brain. For our analysis, we focussed on the motor cortex (M1C) and the middle temporal gyrus (MTG) - two of the profiled regions that had a large number of nuclei and are also largely overlapping in terms of their milieu of cell subsets. Therefore, we designated the nuclei of the M1C region as the reference data set and the nuclei of MTG as the query data set.

Another important feature of this data set is that the annotations in the original study were done at different levels of granularity. Here, we consider two of these - a coarse level with 22 labels and a fine level (nested in the coarse one) with 214 labels (Suppl. Figure 8A). This will allow us to evaluate how the consensus scores of popV can help determine the desirable level of prediction granularity, balancing between resolution (more granular is better) and accuracy (more granular is more difficult).

Considering integration-based algorithms in popV, we find that our query and reference data sets are well mixed, which agrees with our prior expectation on the similarities between the two respective brain regions (e.g., scANVI integration in Figure 3A). Taking into account the coarse-grained annotation level, we find that the most prevalent labels (represented by more than ten nuclei) were correctly transferred using popV (Figure 3B and Suppl. Figure 9). The two exceptions are also highlighted as low consensus (Figure 3B, C). In the first case, four of the algorithms in popV assigned a label of *Upper rhombic lip neurons* to a small group of nuclei in the query data set, although it is not expected that cells of this type will be found in the brain cortex. The remaining three algorithms (here OnClass was not used as the labels were not matched to the cell ontology database) assigned these nuclei with a label of *Upper-layer intratelencephalic neurons* and *deep-layer intratelencephalic neurons*, which matches their original (hidden) label. This case demonstrates that manual scrutiny of the predictions made by the different algorithms in low-consensus areas can help identify and resolve issues with automated annotations (in this case, using prior knowledge on brain physiology). In the second case, we once again find a disagreement between the different algorithms, this time for a group of nascent oligodendrocytes. While four algorithms in popV assign to most of these nuclei a label of *Commited Oligodendrocytes precursors*, the remaining three choose *Oligodendrocyte precursor*. Similarly to the previous case, the minority vote of popV matches with the original (hidden) reference annotation, and highlighting this region as low consensus can help recover the correct label by manual inspection of the different predictions.

**Figure 3:**
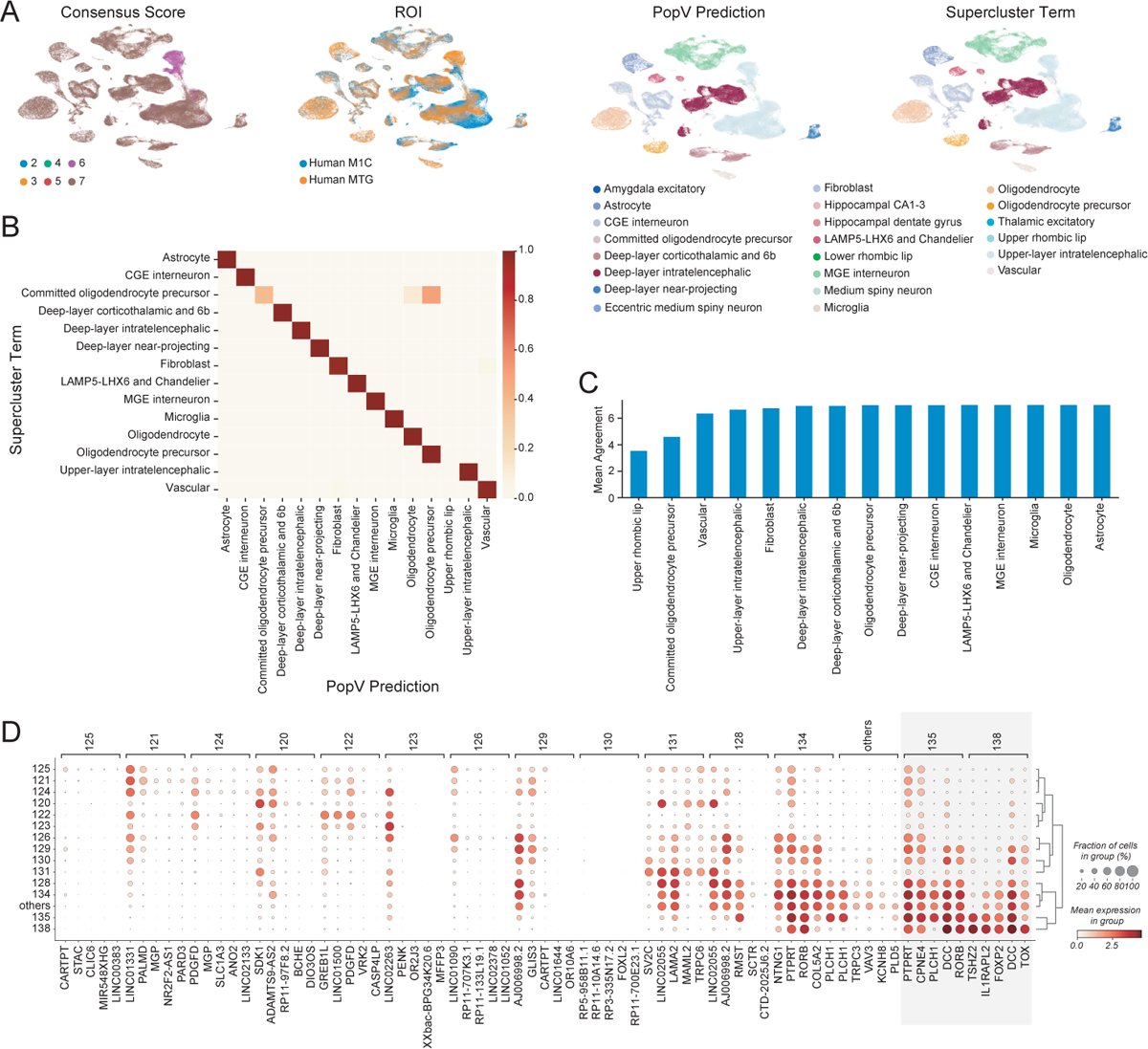
PopV accurately performs labeling of cell types across different brain regions and highlights region-specific neurons. **A**, UMAP embedding after scANVI integration of reference nuclei (motor cortex, M1G) and query nuclei (medial temporal gyrus, MTG) labeled with consensus score, brain region (ROI), popV prediction, and original annotation (supercluster term). **B**, confusion matrix for the cell types predicted by popV and their respective manual annotations highlights the agreement between both annotations. **C**, mean agreement score per cell type shows that confused cell types also exhibit a lower agreement score and can be detected based on their score. **D**, differentially expressed genes for cluster ID for *upper-layer intratelencephalic neurons*. Highlighted are cluster IDs 135 and 138 which are over-represented in the MTG over the M1G. These clusters show an overexpression of FOXP3 and TSHZ2 and are very similar to each other.

Interestingly, the only other coarse-grained label that did not show a perfect consensus is *Upper-layer intratelencephalic neuron*. Here, we find a subset of nuclei for which six algorithms assign the label of *upper-layer intratelencephalic neuron* while scANVI assigns them as *deep-layer intratelencephalic neurons*. Although the (hidden) reference annotation agrees with the majority vote of popV, it is interesting to see a lack of perfect consensus in this case. Indeed, the *upper-layer intratelencephalic neurons* in M1C and MTG clustered separately in the original study, showing that there are region-specific effect on the transcriptomes of these nuclei. To explore this, we calculated differentially expressed genes for all cluster IDs in upper-layer intratelencephalic neurons (query and reference) and found several markers that are specific to those query-specific clusters of *upper-layer intratelencephalic neurons* (clusters 135 and 138 in the original study; highlighted in Figure 3D and Suppl. Figure 8B). Furthermore, we found several genes that distinguish query-specific clusters from reference ones, which are also expressed in deep-layer intratelencephalic neurons (TSHZ2, FOXP2, PDZRN4 and RXFP1), thus providing some reasoning for the lack of consensus from popV. More generally, this case demonstrates that a lower consensus score allows one to highlight subtle differences in cell type phenotypes between query and reference cells and, if necessary, to manually curate those cells.

Next, we wanted to explore which of the granular labels can be reliably refined by applying popV with the fine-grained labels of the reference (M1C) data set. Based on the resulting consensus scores, we found that for some groups of nuclei, a more refined annotation can be reliably achieved, while for others, it was more difficult to go beyond the coarse-grained level of annotation (Suppl. Figure 10). As an example for the first case, we found that the group of *upper-layer intra-telencephalic neurons* in the query (MTG) data set can be further divided into subgroups with a high level of consensus and with a high level of agreement with the (hidden) query labels (Figure 8D). In particular, although in the original study those subsets of *upper-layer intertelencephalic neurons* are associated with distinct labels, there is no discussion about their functional distinction. However, we find that two of these subclusters (clusters 135 and 138 in the original study; highlighted above) are distinguished from all other *upper-layer intertelencephalic neurons* by their expression of FOXP2 (Figure 3D), which is genetically associated with speech disorders in humans [23]. Therefore, this higher resolution annotation points to specific subsets of FOXP2+ neurons in the MTG, a region that has a critical and long-studied role in language processing.

An example for the second case are oligodendrocytes. When attempting to conduct a finer annotation of these cells in the query data, we achieve low consensus scores. When we examine the different subsets of oligodendrocytes in the fine-grained reference, we indeed find little transcriptomic differences, which makes the classification task difficult. This highlights a case where coarse-level annotation is safer, and any refinement of the labels requires manual curation - deciding whether minor differences in transcriptome are indeed relevant for cellular function and justify distinct labels.

### PopV provides useful label transfer in case of drastic differences in cellular composition

After highlighting that popV is capable of detecting query-specific cells and that the consensus score is capable of highlighting these cells, we studied whether this can also be achieved when we have very different query and reference data sets. To this end, we studied the annotation of thymus cells using Tabula sapiens as a reference data set and a second study, which profiled thymi from different age groups (fetal, childhood, adolescence, and adulthood) as query [24]. In particular, the thymus undergoes involution with age, and the adult thymus, which we use here as reference, does not accurately represent the structure and function of the thymus in younger individuals. In particular, we anticipate that the reference sample will not provide ample representation of the developing T cell population, which should be prevalent in our query data.

UMAP embedding of the two harmonized data sets (with scANVI) clearly highlights the subsets of query cells that are represented in both data sets vs. the ones that are query-specific. For most of the common cell types, we found a high consensus between the methods and good agreement between popV prediction and the original (hidden) annotation of the query data set (e.g., accuracy greater than 95% for cells with a popV prediction score of 7 and 8; Figure 4B, Suppl. Figuer 13 and Suppl. Figure 14). In contrast, and as expected by the age of the donors in the Tabula Sapiens project, the compartments of thymocytes and developing T cells are almost absent from the reference data set (Figure 4A). Reassuringly, popV assigned low consensus scores to the majority of cells from both of these compartments, highlighting them for manual annotation (Suppl. Figure 16).

**Figure 4:**
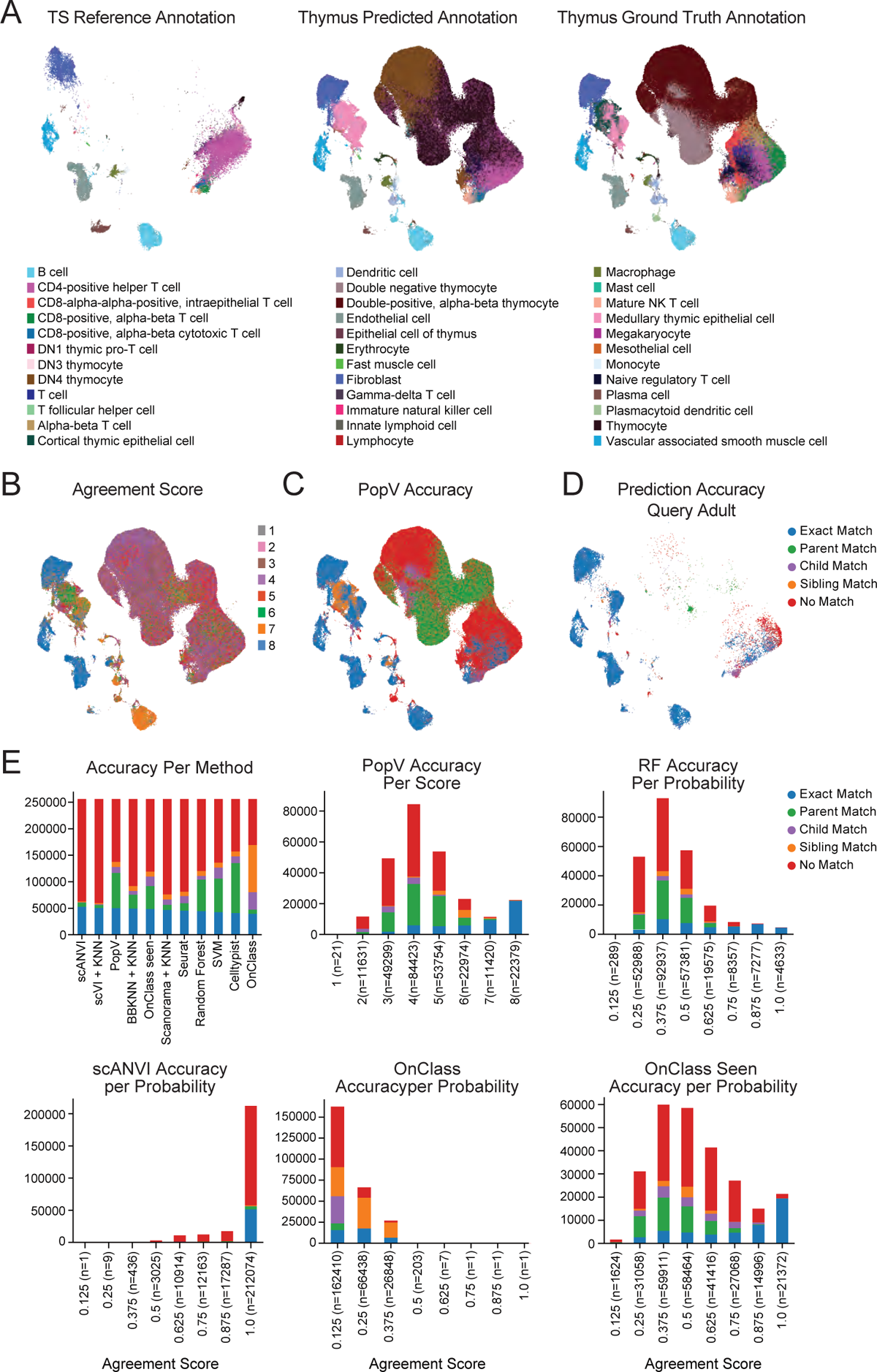
PopV identifies thymocytes as a query-specific cell types and yields highly interpretable consensus scores. **A**, UMAP embedding after scANVI integration of reference cells (Tabula sapiens) and query cells (thymus cells across different age groups) labeled by popV prediction, and original annotation. **B**, popV prediction score overlaid on the UMAP plot. The prediction score is low for thymocytes and higher for most other cell types. **C**, the prediction accuracy of the popV prediction highlights the low accuracy in developing thymocytes. **D**, the prediction accuracy of the popV prediction in adult thymus cells in the query shows high accuracy except for CD8 T cells. **E**, all methods show comparable accuracy with varying numbers of *Parent Match*. PopV, RF, and OnClass_seen show a good correlation of certainty and accuracy, while popV has the highest number of confidently annotated cells.

We identified two other cell populations that are underrepresented in Tabula sapiens compared to the query data set, which are *cortical thymic epithelial cells* (also associated with involution [25]), and *plasmacytoid dendritic cells*. Similarly to our previous examples, we find that the consensus score associated with *cortical epithelial cells* is indeed low, with a variety of annotations assigned to these cells by the different algorithms, including *fibroblasts* and *medullary epithelial cells* (Suppl. Figure 14). The low consensus score suggests that manual curation of this group of cells is needed. In this case, the manual assignment of the correct out-of-reference label is relatively straightforward using PSMB11, an established marker of *cortical thymic epithelial cells* that is not expressed in any cell in the Tabula sapiens reference.

For *plasmacytoid dendritic cells*, all algorithms except SCANORAMA + *k*NN predicted that those cells are *B cells* or *plasma cells*. SCANORAMA + *k*NN predicted that those cells are *dendritic cells*. Even OnClass, which can predict cells not present in the reference data set, predicted those cells as *antibody secreting cells* or *lymphocytes of B lineage* with not a single cell correctly predicted as a *plasmacytoid dendritic cell*. However, these query cells expressed high levels of CLEC4C and IL3RA and were therefore correctly labeled as *plasmacytoid dendritic cells*. As two thirds of *plasmacytoid dendritic cells* have a score of 5 or lower, manual identification of these cells is possible and displaying marker genes identify those confidently wrongly annotated cells, as above.

The only cell fraction that had a high consensus score but low accuracy is a group of cells labeled as *Endothelial cells* by popV, while annotated as *Lymphocytes* in the original (hidden) annotation of the query data set. However, these cells express CAVIN2, TFPI, which fits well with an annotation as endothelial cells. We found that their gene expression aligns well with lymphatic endothelial cells. Therefore, it suggests a wrong annotation in the query data set and a correct prediction by popV.

Overall, this demonstrates that the consensus score yields an interpretable metric for prediction accuracy, and that it helps handle cases of discrepancies between the query and reference data set.

## 3 Discussion

We have developed popV, an ensemble method for cell type annotation, to yield an interpretable certainty quantification for the task of cell type annotation. We have demonstrated throughout this manuscript that in various scenarios with different sequencing technologies, various cell type resolutions, and various overlaps of reference and query data sets, popV yields a confidence score that is well correlated with the actual accuracy of cell type transfer. We demonstrated that the prediction score can predict cell types that are specific to the query data set (MTG-specific neurons), incorrectly annotated in the reference (CD4 T cell subsets in Tabula sapiens) or in the query data set (*lymphatic endothelial cells* in the thymus), or cell types that are not annotated in the reference data set while present in both data set (*lung intermediate capillary endothelial cells* in Tabula sapiens).

PopV is implemented as an easy-to-install, open-source Python tool. The code base is designed so that adding additional cell type classification algorithms is straightforward, thereby allowing researchers to mitigate the risk of choosing a single algorithm (i.e., circumvent the no “one size fits all” problem). We expect future annotation tools to be developed and popV to be used as a tool to handle various biases in these tools and to help quantify certainty in automatic prediction. As an example, upon user request, we included harmony + *k*NN, which was not part of the initial release and therefore not used throughout the manuscript, as a classification model and found popV’s flexible framework to be straightforward in implementing novel predictors.

PopV’s performance is limited by the performance of the underlying predictors. We showed throughout the manuscript that overall popV performed equally well as the single-best method in terms of accuracy. However, the aim of popV is not to improve the accuracy of cell type annotation over the single predictors but to yield a metric of certainty that is easy to interpret and well-calibrated. Indeed, we found that algorithm-intrinsic certainties tend to be poorly correlated with the accuracy of cell type annotation. This is also reflected in a recent study that highlights the low calibration of conventional classification tools [26]. Conversely, we demonstrated that the popV consensus score is highly associated with accuracy and that it helps identify cases where manual involvement is required.

To make popV a valuable resource for the community, we provide a Google Colab notebook with pre-trained models for every tissue in the final Tabula sapiens publication. Moving forward, we expect popV to be integrated with the annotation platform of the Human Cell Atlas. This will allow code-free and interactive browsing of the results such as prediction, popV consensus score, and visualization of marker genes for manual scrutiny.

## Acknowledgements

This publication is part of the Human Cell Atlas www.humancellatlas.org/publications/. We acknowledge the critical feedback provided by the Human Cell Atlas Cell Annotation Platform team. We thank Leah Dorman for extensive benchmarking of popV. We thank the Graphics Department at Weizmann Institute of Science, especially Ishai Sher, for helping with the graphics in this manuscript. We acknowledge members of the Yosef laboratories for general feedback. N. Y. was supported by the Chan Zuckerberg Initiative Essential Open Source Software Cycle 4 grant (EOSS4-0000000121) for scvi-tools. C. E. was supported by the German Research Council (DFG Walter Benjamin Stipendium 448802458). A.S. is a Chan Zuckerberg Biohub Investigator.

## Author contributions

C.E. and G.X. contributed equally. C.X, G.X., and N.Y. conceptualized the study. G.X. conceptualized the statistical model with contributions from C.X., C.E., M.J., and A.M.. C.E. designed and implemented popV with contributions from G.X., C.X.. N.Y., A.P., A.S. supervised the work. C.E. and N.Y. wrote the manuscript.

## Competing interests

N.Y. is an advisor and/or has equity in Cellarity, Celsius Therapeutics, and Rheos Medicine.

## 4 Methods

PopV is a Python package available via the Python Package Index (PyPI). Further details on popV, the source code, and a model tutorial are available at https://github.com/YosefLab/popV.

### Data sets

#### Tabula sapiens

Tabula sapiens was used throughout the manuscript as the reference data set. Tabula sapiens was downloaded from CELLxGENE https://cellxgene.cziscience.com/ collections/e5f58829-1a66-40b5-a624-9046778e74f5. The expression data was set to the raw object of the h5ad object, which contains count data for all cells and genes. This yields 483152 cells and 58559 genes. We filter every cell that has less than 10 cells in a respective tissue as the *k*NN used in popV cannot predict cells with less than 8 examples (15 nearest neighbors by default) (Suppl. Table 1). We confirmed that all cell types are present in the recent version of the Cell Ontology downloaded from https://github.com/obophenotype/cell-ontology/tree/v2023-02-19. Furthermore, we validated that the cell type annotation was not donor-dependent in Tabula sapiens. Tabula sapiens was annotated on a per donor basis and for early donors cell type labels have different names for the same cell type compared to latter donors. To reduce the effect of this inconsistency, we excluded several samples (Suppl. Table 2). Additionally, we found a strong batch effect between some 10X samples. After contacting the original authors, we found the 10X chemistry to be the reason for this and created a new metadata column containing the correct assay. The corrected assay can be accessed through https://doi.org/10.5281/zenodo.7587774. All models were trained using a batch covariate of concatenated donor and assay and separated by tissue (Suppl. Table 3 and 4).

#### Lung Cell Atlas

Data was downloaded from CELLxGENE https://cellxgene.cziscience.com/collections/5d445965-6f1a-4b68-ba3a-b8f765155d3a. We relabeled the cell types to attain conformity with the Cell Ontology (Suppl. Tab 6). Additionally, we filtered all blood samples collected for the construction of the Lung Cell Atlas. We created a concatenated column of sample ID and assay and used these concatenated metadata as the query data set batch key in popV (Suppl. Table5. Throughout this manuscript, the query data set label was not used as input to scANVI as the general application of popV works with an unlabeled query data set. The lung cell atlas contains 75071 cells in total and 39 unique cell types were used as cell ontology labels out of 59 unique cell types in the original lung cell atlas.

#### Brain data set

Data was downloaded from CELLxGENE https://cellxgene.cziscience.com/collections/283d65eb-dd53-496d-adb7-7570c7caa443. We downloaded *Dissection: Cerebral cortex (Cx) - Precentral gyrus (PrCG) - Primary motor cortex - M1C* and *Dissection: Cerebral cortex (Cx) - Middle Temporal Gyrus - MTG* as the two cortical regions with the largest number of cells. Original cell type labels were used for this data set and we used respectively *cluster_id* and *supercluster_term* as the cell type key. We remove cells labeled with the super-cluster terms *Splatter* as well as *Miscelleanous* as these likely contain low-quality cells where manual annotation was failing. For all downstream metrics, we removed cell types with less than 10 cells in each cell type label as we found those to be reflective of nuclei from distinct brain regions (*Medium spiny neuron*, *Hippocampal dentate gyrus*, *Hippocampal CA1-3*, *Amygdala excitatory*) We decided against using the labels that conform to the Cell Ontology, as all neurons were labeled with the same cell ontology term “neuron”, which does not reflect the heterogeneity of these cells. The cell type labels termed *subcluster_id* were the finest level of annotation. However, we found little evidence in the transcriptome of nuclei to trust those labels and were excluding those from analysis.

#### Thymus data set

Data was downloaded from https://cellxgene.cziscience.com/collections/de13e3e2-23b6-40ed-a413-e9e12d7d3910 and the data was analyzed using the same CELLx-GENE access link. We relabeled the cell types to achieve granularity comparable to the reference data set (Suppl. Tab 6). For subset analysis, fetal samples were filtered to every *Development Stage* containing *th week* as a substring, adult samples were filtered to *human early adulthood stage*. We use the donor ID and assay as the query data set batch key in popV (Suppl. Table8. The thymus data set contains 255901 cells in total and 28 unique cell types were used as cell ontologies out of 31 unique cell types in the original thymus data set. All cells in this data set were labeled according to the cell ontology. However, we decided to summarize all CD4-positive, as well as all CD8-positive T cells, into a common cell type to make the annotation granularity comparable between reference and query data sets (Suppl. Figure 12). We additionally summarized all B cells in the query and reference data set to be annotated as B cells, as the label of B cells in Tabula sapiens showed strong donor inconsistencies and summarized all endothelial cells to be labeled as endothelial cells to harmonize the granularity of cell type labels.

#### Model parameters

We use eight different cell type annotation algorithms and in the following explain our parameters for those annotation algorithms as well as the data preprocessing pipeline that we use for popV. For UMAP embedding, we used scanpy default parameters except a *min_dist* of 0.3. For the ‘*k*NN classifier, we use uniform weights and *n_neighbors* equal to 15 in *sklearn.neighbors.KNeighborsClassifier*. The classifier is first trained on all reference cell labels and is then applied to all query cells in prediction mode. To increase performance of this classifier we use an sklearn pipeline and use PyNNDescent for neighbor computation [27]. All default parameters for the underlying methods can be changed using a dictionary *method_kwargs* upon calling *popv.annotate_data*

#### Pre-processing

Every data set was pre-processed using the *Process_Query* function in popV. The input parameters of *Process_Query* are explained in the popV documentation. If using a pre-trained model folder, both reference and query data sets are subsets of the same genes. It is validated that both data sets contain raw counts. The cell type labels in the reference data set are subsampled to 300 labeled cells by default to reduce the runtime of the underlying methods. The intersection of genes between the query data set and the reference data set is taken, and both data sets are concatenated. We remove all batches in the query and the reference data set that contain less than 9 cells in total as otherwise BBKNN is failing and remove all cells with less than 20 counts. Highly variable genes are computed using *seurat_v3* flavor in scanpy [28] and by default 4000 genes are selected. Count data is stored and counts are further normalized to *10k* counts and the *log1p* function is applied for methods that require normalized data and stored in a separate layer. For the computation of principal components, the count data is scaled to unit variance. These principal components are used for SCANORAMA and BBKNN. All keys used to set up the model are stored in the *uns* field of the anndata object [29]. PopV has three different modes:

- retrain: Trains all methods from scratch and stores the classifier to reuse them on other data sets. This hugely benefits from a GPU to train the scVI and scANVI algorithms as well as the OnClass algorithm.
- inference: Uses pre-trained methods to classify query and reference cells. It computes a joint UMAP embedding of query and reference cells and by default uses all eight methods. Trains scVI and scANVI models for 20 epochs using scArches query embedding [19].
- fast: Uses pre-trained methods to classify only query cells. Computes a UMAP embedding of query cells if enabled. Skips Scanorama and BBKNN data integration as those recompute an embedding instead of projecting cells into an existing embedding. Trains scVI and scANVI models for 1 epoch using scArches query embedding.

#### BBKNN

Batch-balanced K nearest neighbor is a data integration method. To integrate the data sets, BBKNN takes the nearest neighbors from each batch to construct a balanced neighborhood graph. This nearest-neighbor graph can then be used as a batch-corrected graph embedding of the data [13]. The default settings for popV and those used throughout the manuscript are 50 principal components, 8 *neighbors_within_batch*, and the *angular* metric. We found that the angular metric outperforms a standard Euclidean metric in our use case. We use the implementation of BBKNN in *scanpy.external.pp.bbknn*. The batch-balanced nearest neighbors are used as a precomputed metric in *sklearn.neighbors.KNeighborsClassifier* and used as input for UMAP dimensionality reduction.

#### SCANORAMA

SCANORAMA is a data integration method. SCANORAMA searches for the mutual nearest neighbors across data sets and uses panoramic stitching. Cells are then integrated in PCA space using those mutual neighbors. By default in popV and throughout the manuscript 50 principal components are used. We compute a new joint embedding of the query and the reference data set using *scanorama.integrate_scanpy* function. This joint embedding is used for the ‘*k*NN classification and UMAP embedding.

#### scVI

ScVI is a variational auto-encoder that incorporates batch keys as latent variables and provides data integration in its latent space. We use the following non-default parameters for scVI *dropout_rate*= 0.05, *n_layers*=3, *n_latent*=20, *gene_likelihood*=*nb*, *encode_covariates*=*True* and *use_layer_norm*=*both*. The reason for these non-standard parameters is to facilitate integration of a query data set using scArches. For the training parameters, we use by default scVI with *n_epochs_kl_warmup*=20 epochs. We compute the joint latent representation of query and reference data and this joint embedding is used for the ‘*k*NN classification and UMAP embedding.

#### scANVI

In addition to scVI, a classifier is trained during training of the auto-encoder on the positions in latent space to classify cells into the provided reference cell type labels. We continue training based on the trained scVI model to reduce the overall training time. For the classifier in scANVI we use *n_layers*=3 and *dropout_rate*=0.1. Subsampled labels are used as discussed above. We use as training parameters *batch_size*=512 and *n_samples_per_label*=20 to stabilize training of the classifier. Subsequently, the built-in classifier is used to predict cell type labels in the query data set.

#### Random Forest

Random Forest uses an ensemble of classification trees together with random feature sub-setting to regularize the classification trees. The final prediction is the majority vote across the tree ensemble. We use normalized counts (see above) as input for Random Forest and *sklearn.ensemble.RandomForestClassifier* as the classifier. We use non-default parameters as *max_features*=200 and *class_weight*=*balanced_subsample* as we found the best performance using this parameter combination. For training the classifier, subsampled cell type labels are used as described above, as this improves prediction speed.

#### Support Vector Machine

Support Vector Machines find the hyperplane that best separates the data. We use *sklearn.svm.LinearSVC* as the classifier. We use non-default parameters as *C*=1 and *max_iter*=5000 and *class_weight*=*balanced* as we found the best performance using this combination of parameters. For training the classifier, subsampled cell type labels are used as described above, as this improves prediction speed. To allow computation of prediction probabilities, we use *sklearn.calibration.CalibratedClassifierCV*.

#### Celltypist

Celltypist uses a logistic regression framework. We use non-default parameters *check_expression* = *False* and *max_iter*=500 to allow for faster model training. During *celltypist.annotate*, we use *majority_voting*=*True*. As intrinsic probabilities we use *predictions.probability_matrix* as the majority voting purity and not the initial logistic regression probabilities. This is similar to the probabilities used in the Celltypist tutorials.

#### OnClass

OnClass first computes an embedding of the Cell Ontology using Natural Language Processing on the cell type names and then applies random walks. This can be embedded using Singular Value Decomposition [30]. Then a bipartite neural network was optimized to allow classification of the reference cells. This network is then applied to unannotated cells. By design, this allows classification of unseen cell types in the cell ontology term-based low-dimensional embedding. We downloaded the obo Ontology files in version *releases/2023-01-09*. To allow fast retraining of sentence embedding, we use *sentencetransformer.SentenceTransformer(’all-mpnet-base-v2’)* as the NLP model. This is a newer NLP model than in the original OnClass publication, but allows for faster convergence. We encode all descriptions or cell type labels in the obo file. We provide notebooks to retrain with newer releases of ontology files or different species. We provide several ontology files in our Github repository, these are *cl.obo* which is the downloaded file from https://github.com/obophenotype/cell-ontology. *Cl.ontology* which is a file containing only the *is_a* cell type relationships from the *cl.obo* file and *cl.ontology.nlp.emb*, which contains the embeddings of the *cl.obo* file. As count data, we use normalized data (see above) and disable the options to recompute this normalization in OnClass. OnClass provides the option to use batch integration using SCANORAMA. We disabled this option to not bias the prediction based on the performance of SCANORAMA. We found no sign of a strong batch effect in cell type prediction. For OnClass, we provide two different cell type labels. OnClass_seen is the prediction of all cells limited to the cell types in the reference data set, while OnClass_prediction contains for each cell type the final output of the OnClass model. OnClass currently only outputs predictions after step 2 for 10% of the cells, even though it computes those on all cells. We found this procedure wasteful and implemented our own version that outputs labels after step 2 on all cells.

#### Harmony

Harmony was not used throughout the manuscript. However, we found it to perform better on large scale data set (more than a million cells and 50 batches) than SCANORAMA. Harmony uses soft-Kmeans clustering and shifts the centroids of those Kmeans clusters to allow for batch correction [17]. We use 50 principal components as default input to harmony. We use the efficient GPU enabled version of harmony https://github.com/lilab-bcb/harmony-pytorch. We compute a new joint embedding of the query and the reference data set using *harmony.harmonize* function. This joint embedding is used for the ‘*k*NN classification and UMAP embedding.

#### Seurat

Seurat uses canonical component analysis and nearest neighbor matching to integrate first the reference data set and then maps the query data set to this integrated data set [21]. Seurat is not part of popV as copying a data frame to R can be a time consuming step for large data. However, we compared the performance of popV with Seurat for the Lung Cell Atlas. We computed 2000 genes using *FindVariableFeatures* with *vst* transformation. We called *FindIntegrationAnchors* using 30 CCA components. Aferwards we scaled the corrected counts and computed 30 PCA components. Those components were used in *FindTransferAnchors* and afterwards *TransferData* is called to transfer the labels from the reference data set to the query data set.

#### Consensus voting

As described in the manuscript, we tried majority voting as well as cell ontology-based aggregation of OnClass results. We found the cell-ontology-based aggregation to outperform majority voting 1. For this voting strategy, we take the majority vote (counting all predictions) for all predictors except OnClass. For OnClass prediction, we use the predicted cell type and in addition every cell type along the path from this cell type to the root node of the Cell Ontology graph and increase the score of those ancestors by 1. We take as consensus score the score at each cell type level node and take as popV prediction the cell type node with the highest score. For majority voting, we use the prediction in OnClass_seen and count the predictors who agree on a certain cell type node. The node with the majority of votes is used as the majority voted cell type label. If there is a tie between two nodes in the amount of votes, we use the cell type label that is further down the cell ontology tree meaning the more granular cell type, if there is still disagreement, we use the cell type that is latter in the alphabet to have a deterministic mapping and not rely on the order of cell types in the prediction matrix.

### 4.1 Evaluation metrics

All code for creating evaluation plots is available in the popV package as a *_reproducibility* module. We will discuss those metrics here. For displaying the translations of cell type terms we use alluvial plots that highlight the corresponding cell types before and after translation to a Cell Ontology conform term.

#### Accuracy metrics

If no cell ontology graph is available, we use F1 metrics to quantify accuracy. The micro F1 accuracy computes the amount of exact matches across the whole data set and is a global metric, whereas the macro F1 score computes a per cell type accuracy and averages this across all cell types. The macro F1 accuracy therefore better represents the performance across rare cell types. We found agreement of both metrics in their evaluation of performance but the macro F1 accuracy to be more sensitive as rare cell types are harder to predict.

If a cell-ontology is available we reasoned that the performance of a predictor is preferable if it predicts a closely related cell type, we therefore computed different matching scores. We computed an exact match similar to the F1 score as cell types that are correctly predicted. Parent match means that the predicted term is a node that is one step closer to the root of the Cell Ontology graph, while child match means that the predicted term is one step further away from root. Sibling match means that the cell type is two steps away from the correct cell and has the same depth as the original cell type in the Cell Ontology tree. We also experimented with more fine-grained metrics quantifying the distance in the Cell Ontology tree between two cell types but found after manually checking the corresponding cell types those nearest matches to be the correct metric to evaluate classification as cell types that are further away have less similarity.

#### Confusion matrix

We use scikit-learn.metric.confusion_matrix and normalize those entries. We compute these entries for all algorithms and the ground-truth label but also between the different algorithms and the consensus label.

#### Differential expression analysis

We use *scanpy.tl.rank_genes_groups* with default parameters to yield differentially expressed genes and *scanpy.pl.rank_genes_groups_dotplot* to plot those results.

#### Code availability

The code to reproduce the experiments of this manuscript will be made available at https://github.com/YosefLab/popv-reproducibility upon final acceptance of the manuscript. The popV package can be found on GitHub at https://github.com/YosefLab/popV. Documentation and tutorials can be found at https://github.com/YosefLab/popV.

## Supplementary Figures

**Supplementary Figure 1:**
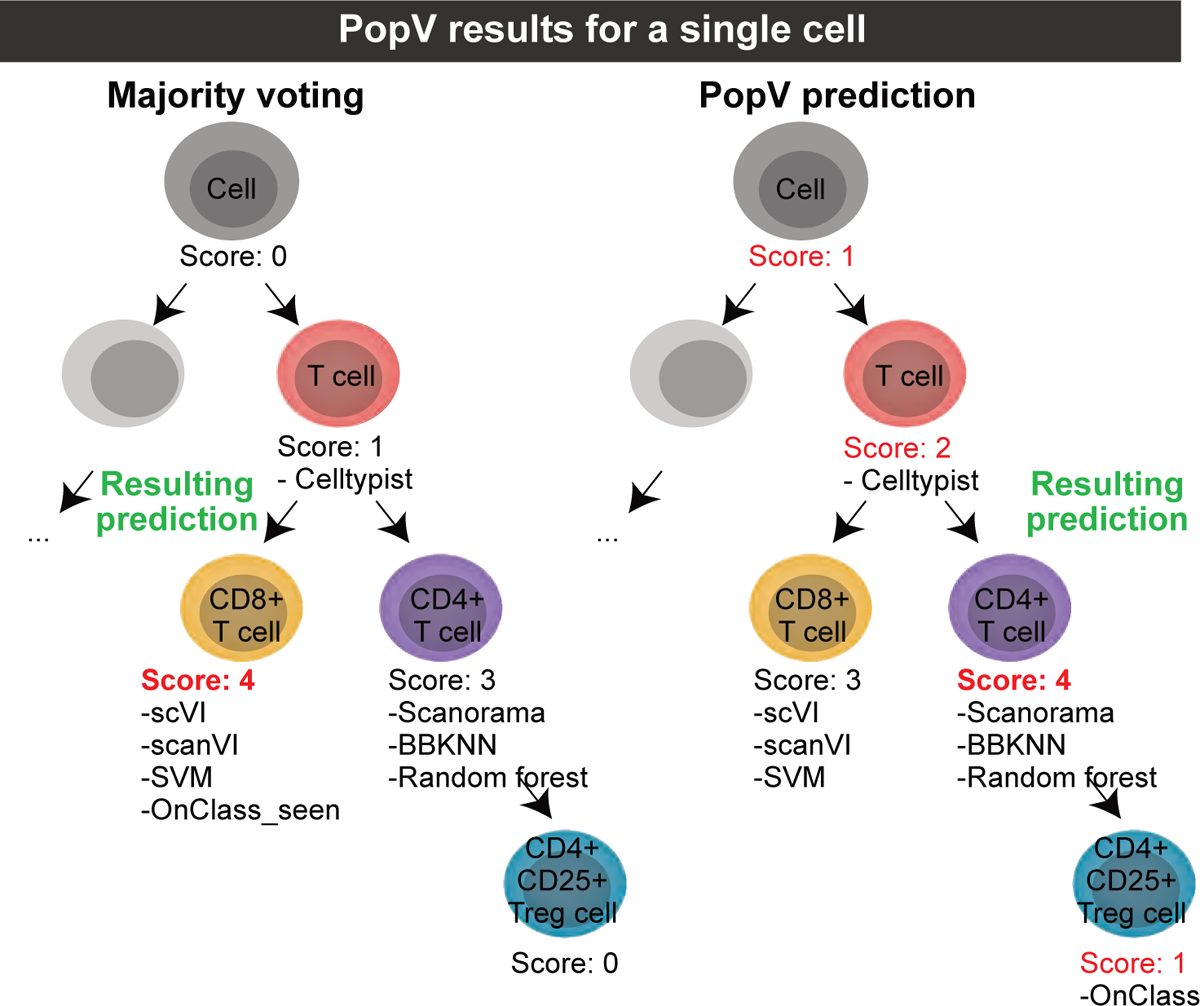
Comparison of majority voting and PopV prediction score. A single cell is annotated by seven different algorithms that are restricted to the cell type labels in the reference data set. For simple majority voting, we use the prediction of OnClass at step 1, where it is bound to seen cell types (OnClass_seen) and count the predictions of each algorithm. In step 2, however, OnClass predicts unseen cell types. Those predictions can have a finer or coarser level along the Cell Ontology hierarchy. A coarser level annotation is called a misclassified cell. To also use finer-level annotations, every cell along the path from the root term to the predicted term receives a score of 1 and the majority voting is performed to receive the consensus cell.

**Supplementary Figure 2:**
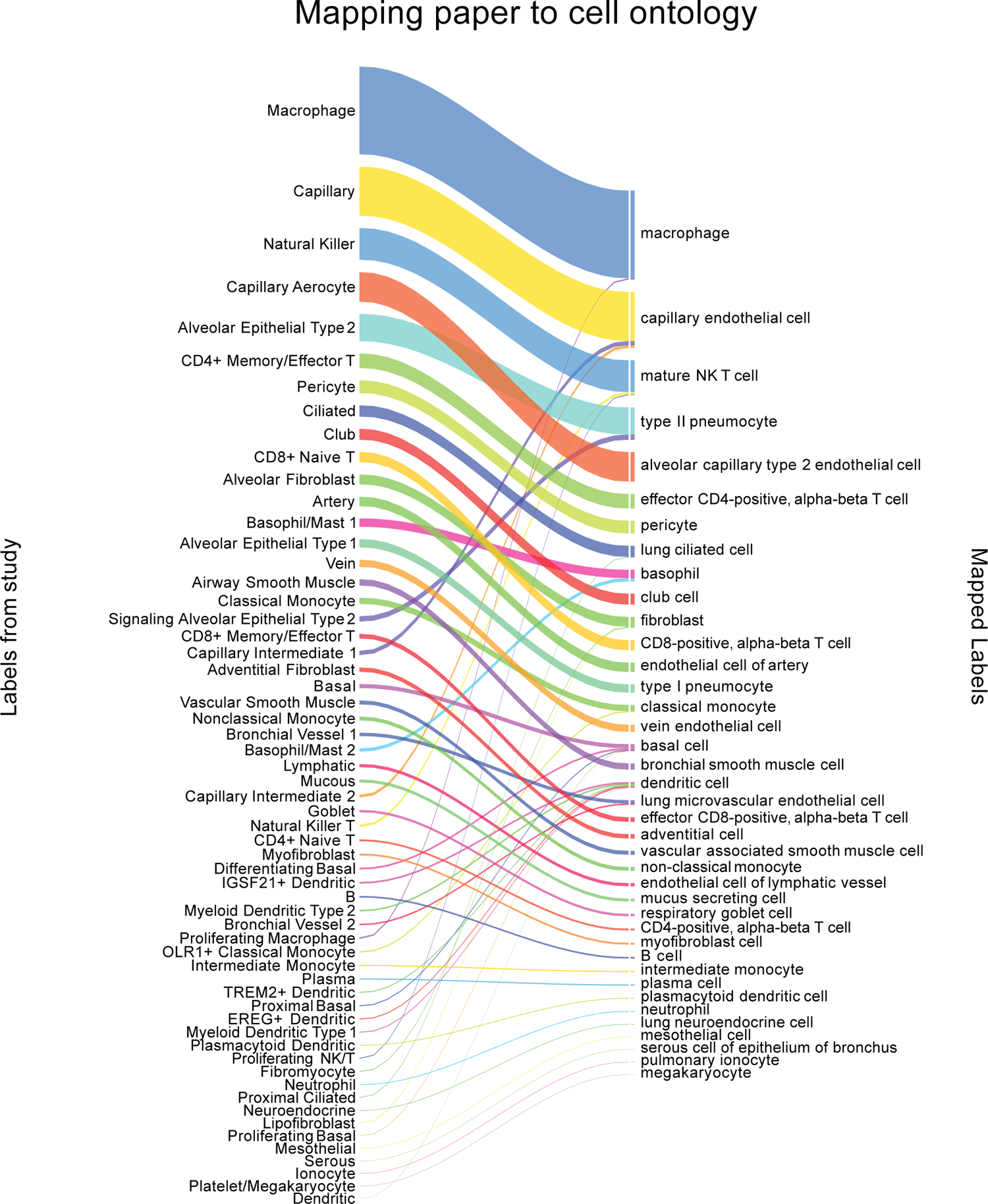
Translation of cell-types in original publication to used cell ontology for lung cell atlas. Cell types were relabeled based on this alluvial plot to assign them to cell ontologies. The closest match in the ontology was manually identified and used as the cell ontology term for every cell type. Of note, several finely resolved cell types were concatenated (e.g. Signaling Alveolar Epithelial Type 2 was renamed to type 2 pneumocyte).

**Supplementary Figure 3:**
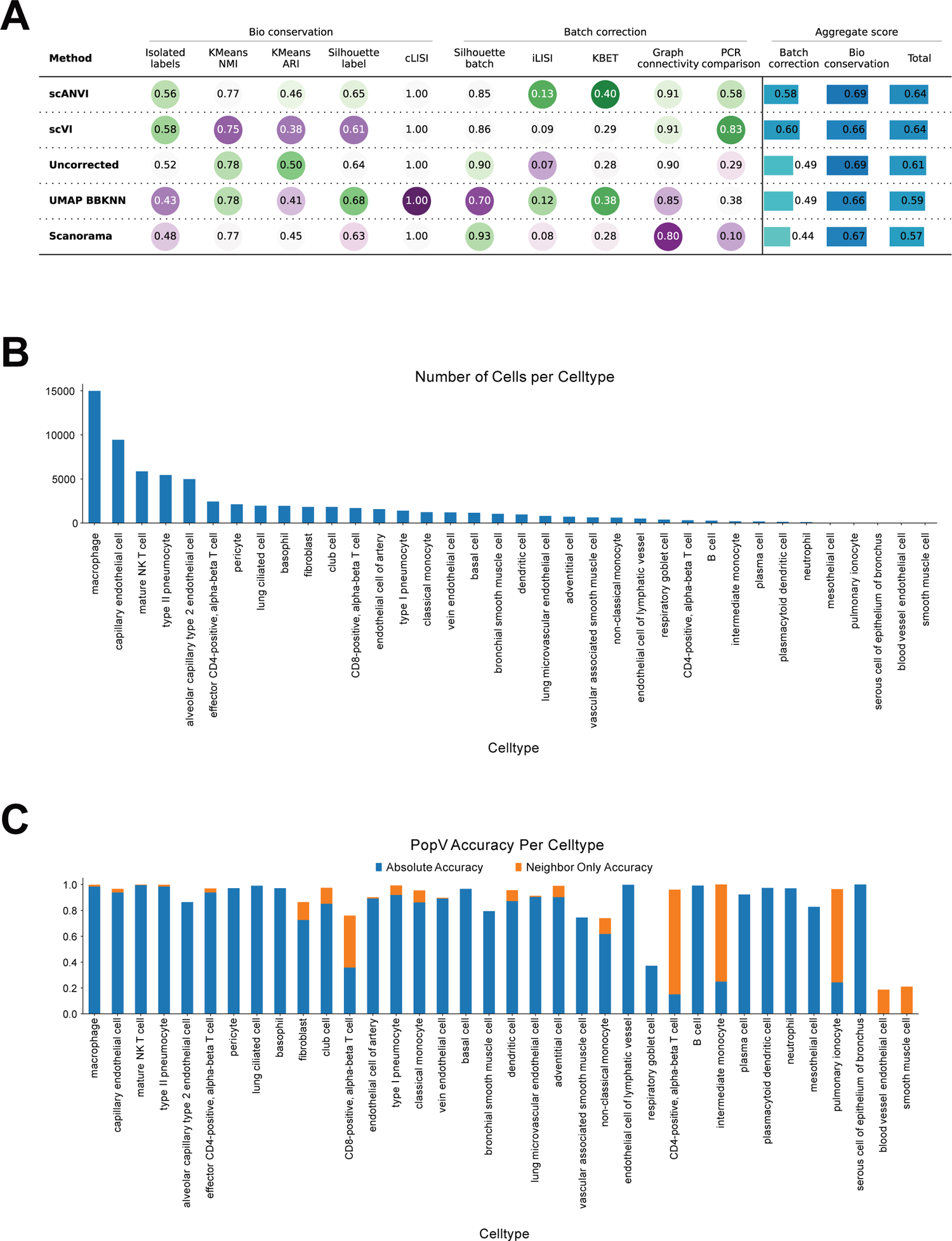
scANVI shows the highest integration of query cells and popV shows low confidence for lowly abundant cell types. **A** scIB metrics comparing integration scores after integrating query and reference data set showed the best integration using scANVI and improvement over uncorrected data. Labels from the original Lung Cell Atlas paper were used to compute cell type-dependent scores and scores were computed only on query cells. **B** Displayed is the number of each predicted cell type in query cells and the accuracy for each annotated cell type **C.** Of note, cells that were rarely predicted (smooth muscle cells and blood vessel endothelial cells showed the lowest accuracy). Most cell types have an accuracy above 0.9.

**Supplementary Figure 4:**
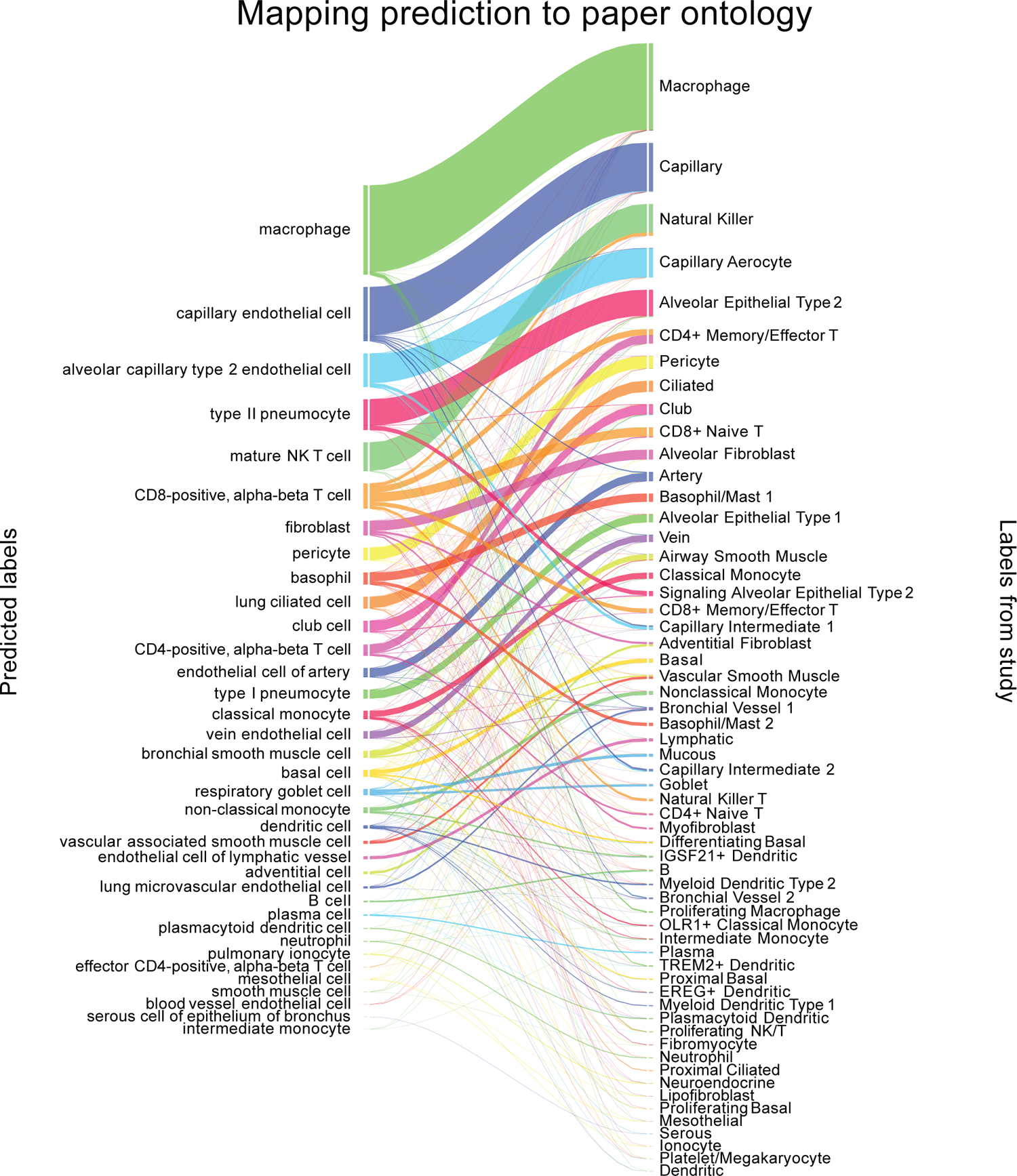
Alluvial plot of annotation in Lung Cell Atlas highlights cells with high diversity of ground-truth labels. The alluvial plot highlights popV predictions of query cells from the Lung Cell Atlas compared to the original annotation. Predicted CD8-positive, alpha-beta T cells e.g. have a wide of original cell-type labels, highlighting the problems with correctly annotating T cells detailed in the main text.

**Supplementary Figure 5:**
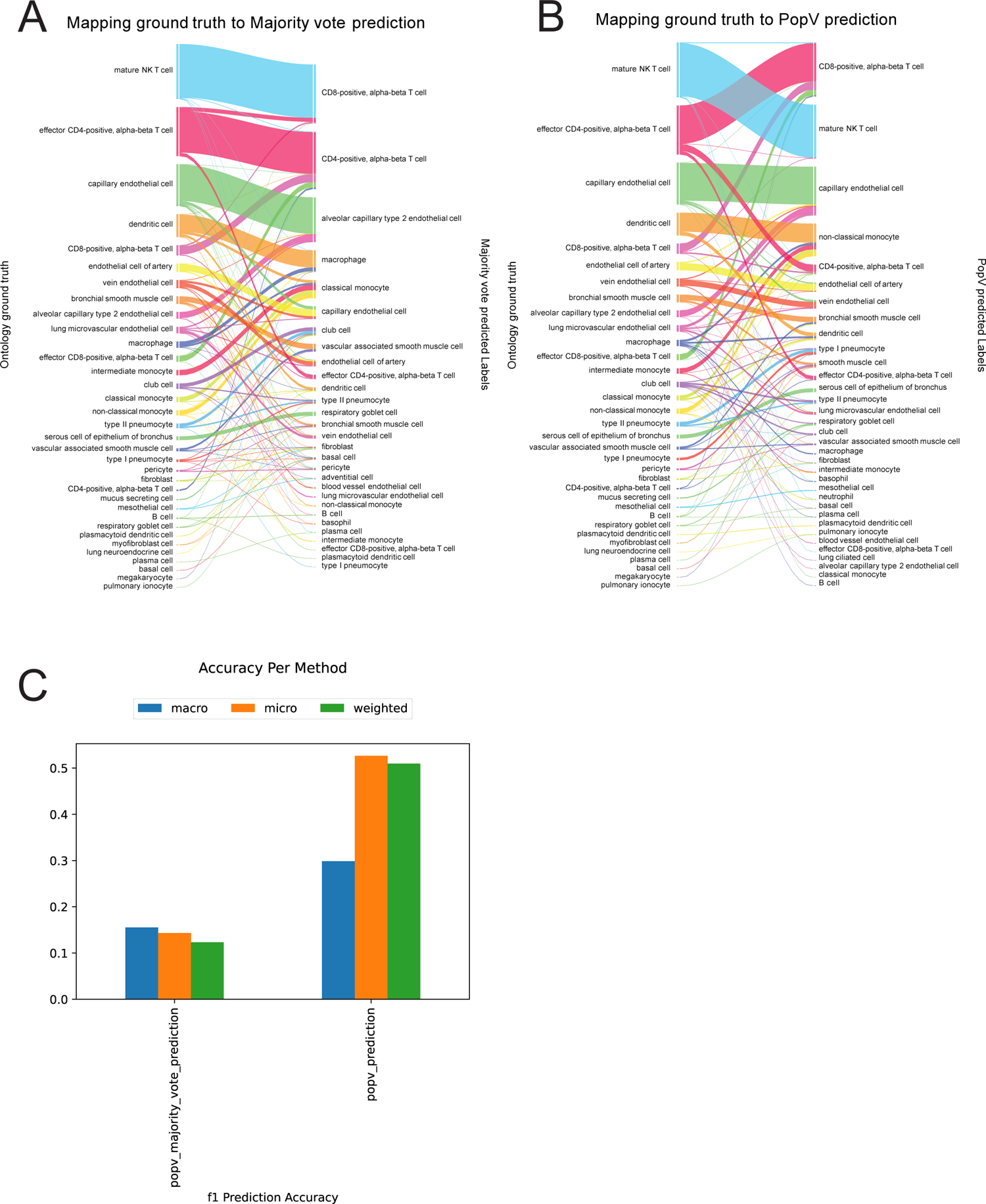
Comparison of consensus scoring and majority voting. Suppl. Figure 1 highlights both scoring options in PopV (majority voting and consensus score). We highlight here cells with different predictions based on both scores. A. Alluvial plots between popV prediction with majority voting and ground-truth labels on these cells. B. Alluvial plots between popV prediction with consensus scoring and ground-truth labels on these cells. C. Comparison of various accuracy metrics between predictions from both scores highlights improved prediction with consensus scoring (popv_prediction) over majority voting.

**Supplementary Figure 6:**
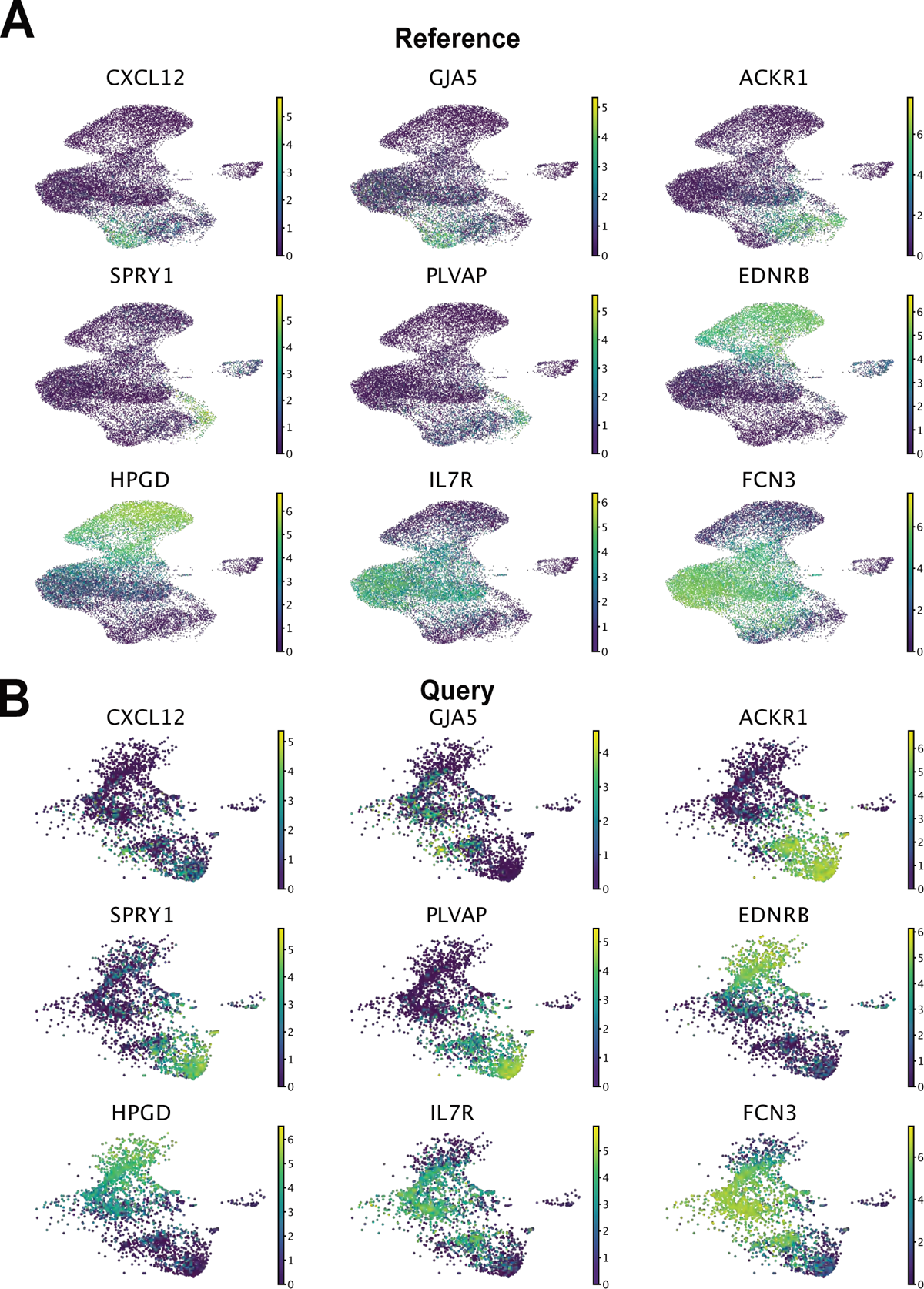
Marker genes for endothelial cells in query and reference cells. We display here marker genes for endothelial cells in both query (A) and reference (B) data set. CXCL12 and GJA are canonical markers for arterial endothelial cells. ACKR1 is a marker for venous endothelial cells. SPRY1 and PLVAP were used as markers for bronchial endothelial cells. EDNRB and HPGD are markers for aerocytes. IL7R and FCN3 are markers of capillary endothelial cells.

**Supplementary Figure 7:**
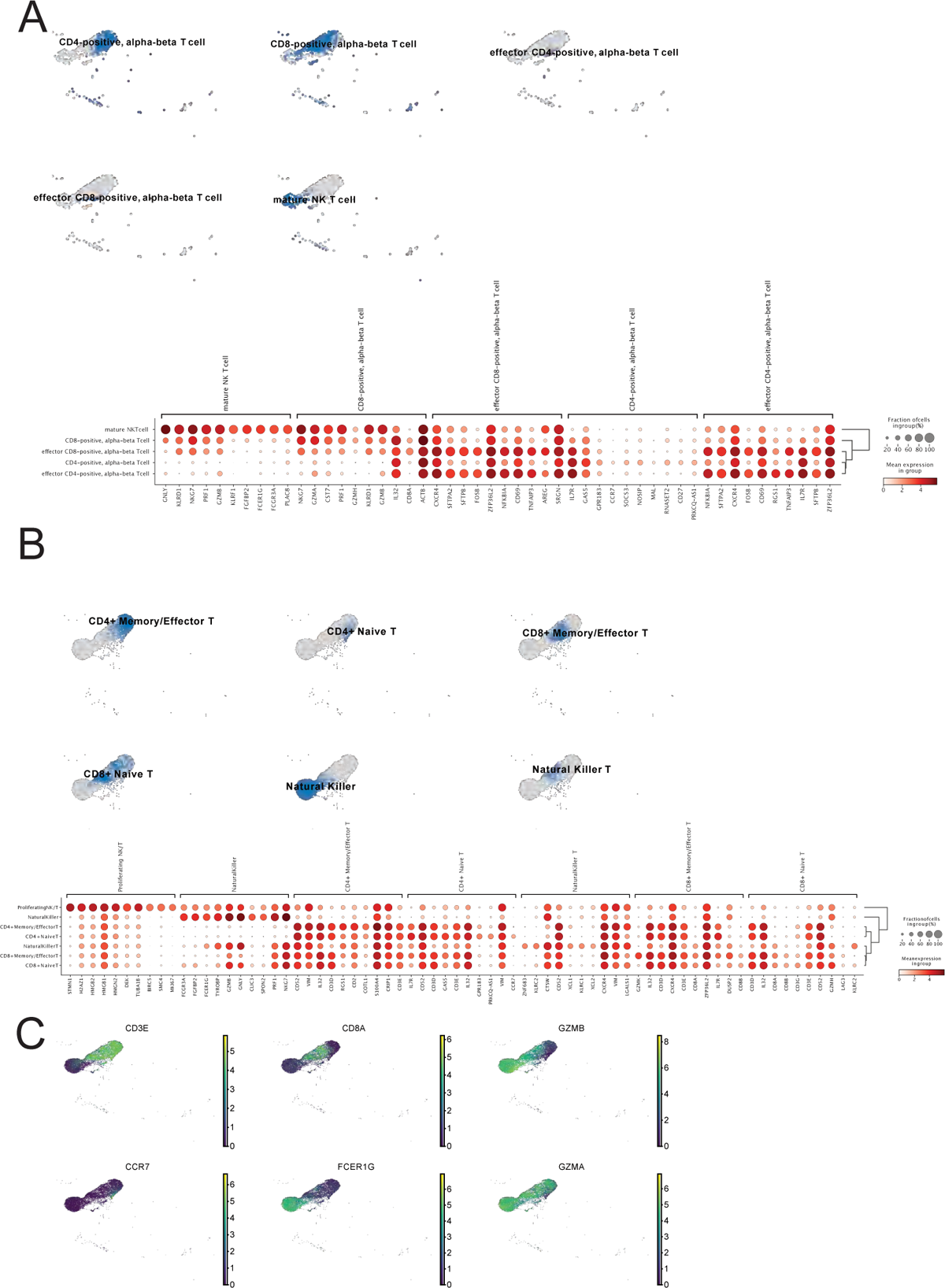
Analysis of T cell sub-clustering in Lung Cell Atlas and Tabula sapiens. **A**. UMAP of cells from the reference data set labelled by cell-type labels from Tabula sapiens, highlights overlap of those labels in integrated space with no clear distinction between CD4 and CD8 T cells. Differential expression analysis identifies surfactant protein genes as markers for annotated effector T cells, which is due to ambient counts and no strong marker gene expression in CD4 T cells. **B**. UMAP of cells from the query data set labelled by cell-type labels from the Lung Cell Atlas shows clear distinction between different cell-types. Differentially expressed genes for those cell types align well with the respective literature. **C**. Canonical marker genes for various sub-types shows a clear split between T-cells and NK cells as well as CD8 and CD4 T-cells. GZMA but not GZMB production is also present in CD4 T-cells.

**Supplementary Figure 8:**
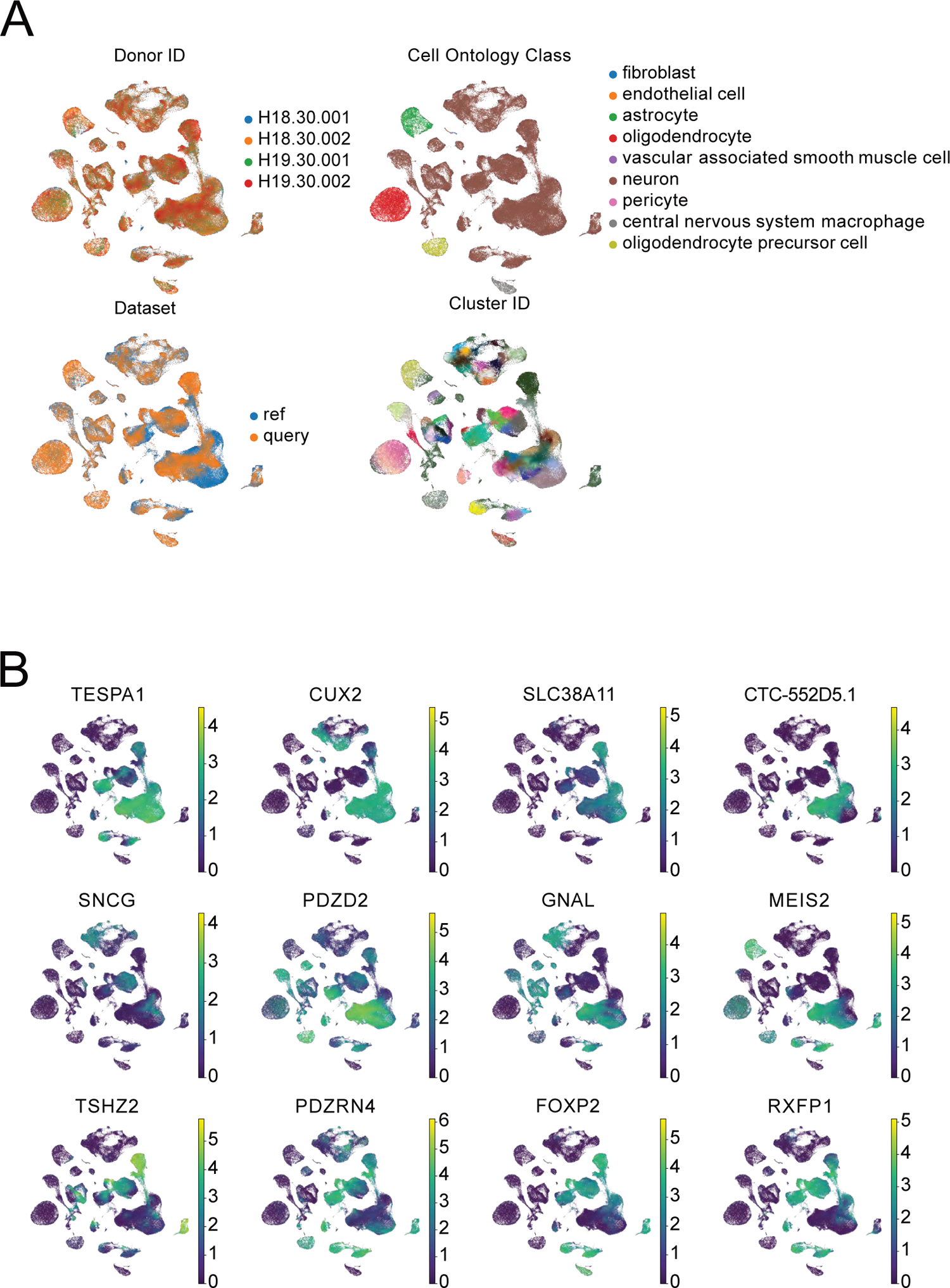
Marker genes show a pronounced heterogeneity in deep-layer inter-telencephalic neurons. Additional meta-data for embedding in Fig 3 3 is displayed in **A**. Data from different donors is well integrated using scANVI embedded space. The cell ontology labels from the paper have a very low resolution and were not used here. Reference and query cells are well integrated. Cluster ID, which is a higher-resolution clustering is conserved in UMAP embedding (legend omitted here to increase readability). **B** We identified marker genes for various cluster IDs in *upper-layer intratelencephalic neurons*. Their expression is displayed here. The first row shows differentially expressed genes for all *upper-layer intratelencephalic neurons*. The second row shows those genes that are more enriched in the M1C. The third row shows genes that are more enriched in the MTG. Genes from MTG are also expressed in *deep-layer intratelencephalic neurons*

**Supplementary Figure 9:**
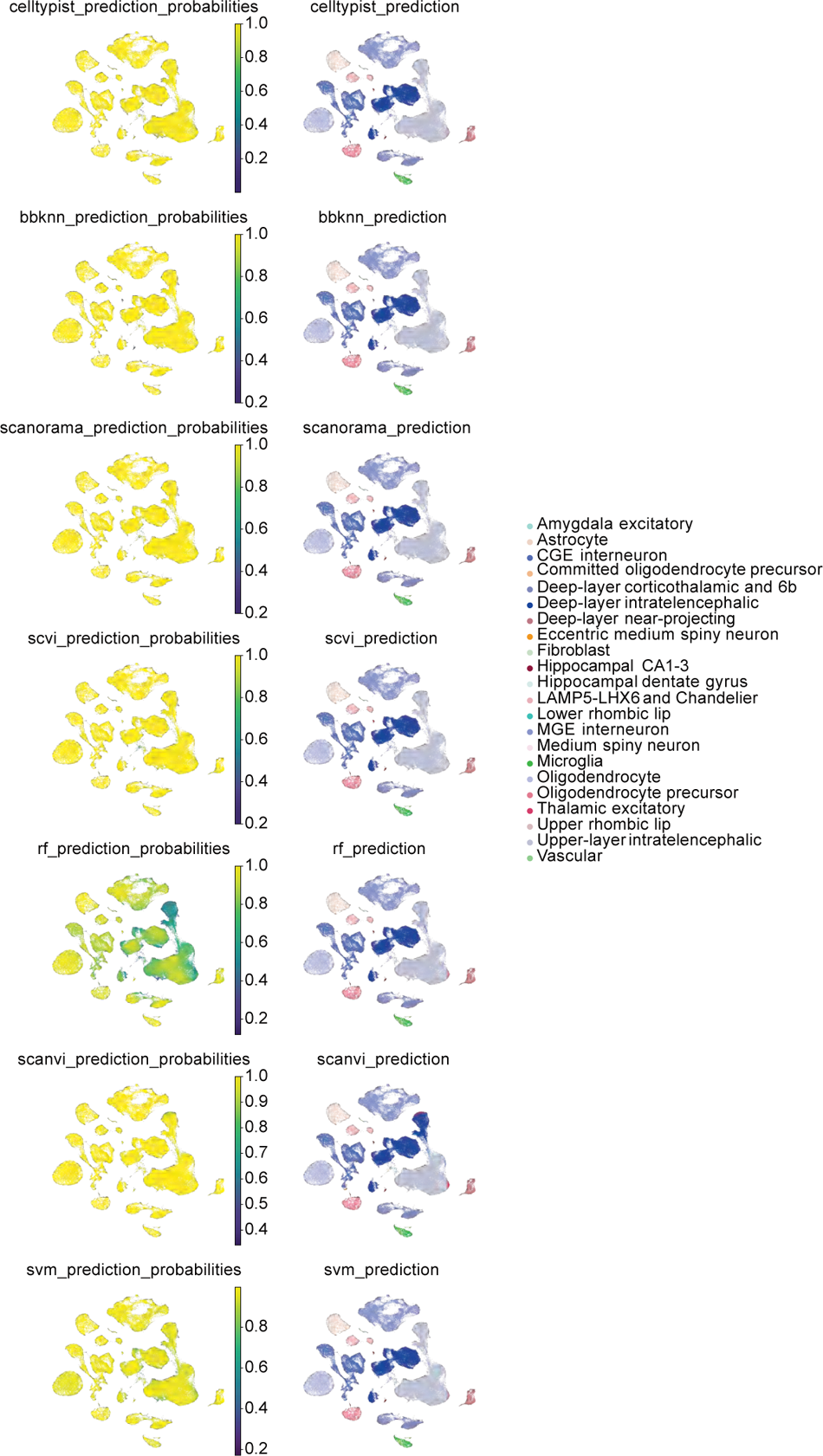
Prediction and certainty for every predictor in brain data. We compared the performance of the various predictors. Except for Random Forest, all probabilities have great certainty in their predictions. Random Forest highlights a more diverse probability across the different cells, highlighting the query specific nuclei described in the main text. Despite its high certainty, scANVI predicts a region of *deep-layer intra-telenchephalic neurons* as *upper-layer intra-telenchephalic neurons*.

**Supplementary Figure 10:**
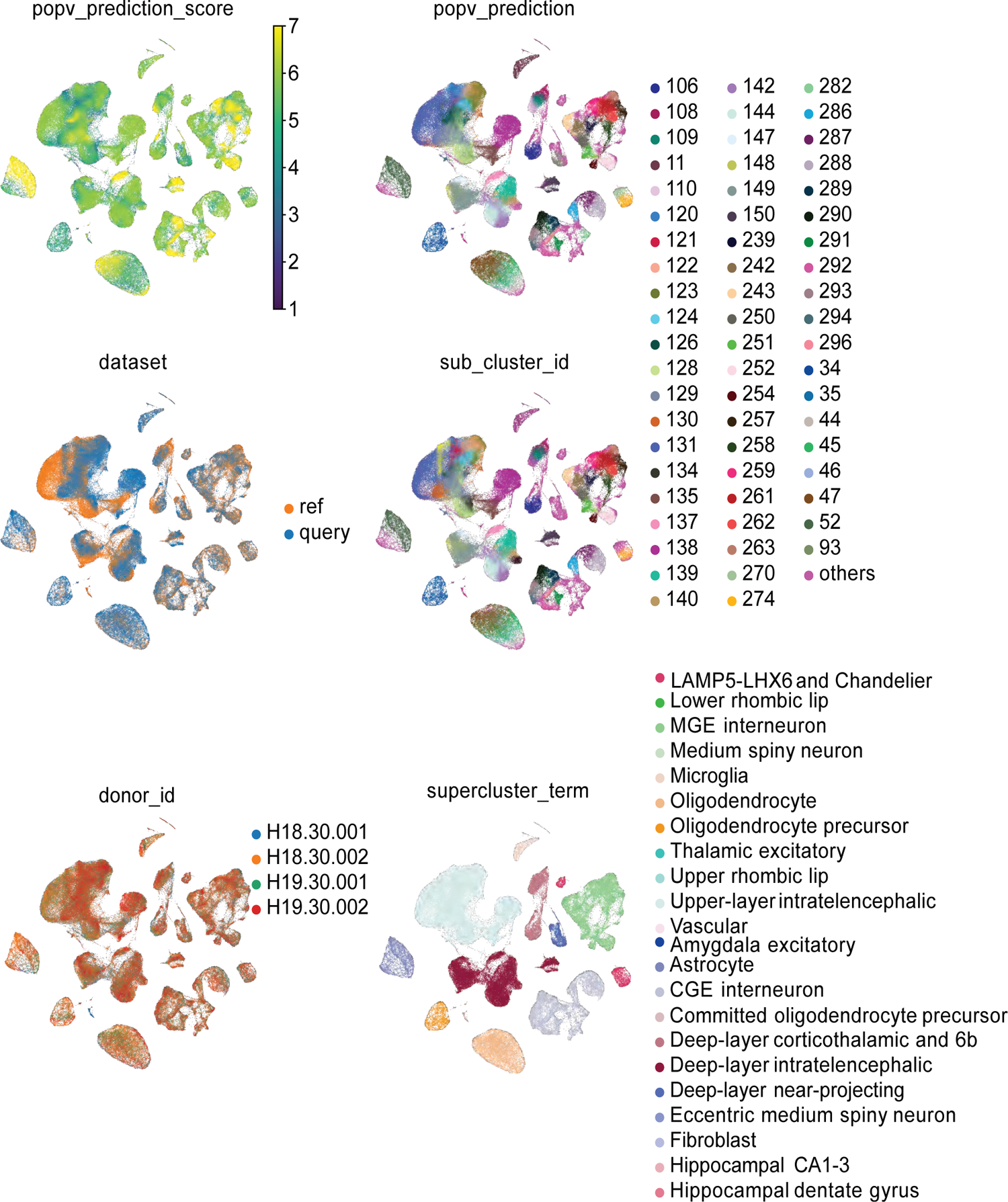
Prediction of cluster ID in brain data using PopV shows low consensus score. In addition to running PopV on the more coarse cell-type labels, we also executed it on the more fine-grained cluster ID. Displayed is the UMAP based on scANVI embedded data using cluster ID as the reference cell-type annotation. The consensus score is clearly lower. Reference and query as well as donors are well integrated. The supercluster term is well conserved in this embedding.

**Supplementary Figure 11:**
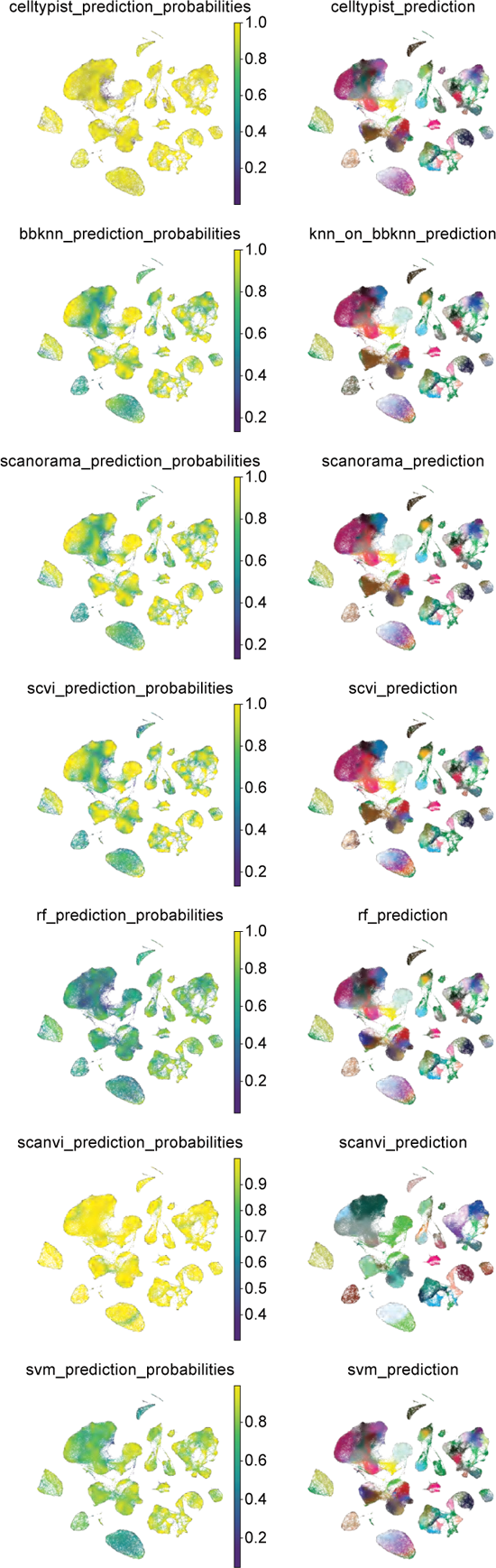
Prediction and probability for every predictor in brain data using cluster ID labels. Due to the higher cell type granularity, the prediction probabilities are much lower when using cluster ID as the ground truth label. Especially for oligodendrocytes every predictor except Celltypist and scANVI display a high uncertainty. Celltypist and scANVI show a high certainty across all cell types.

**Supplementary Figure 12:**
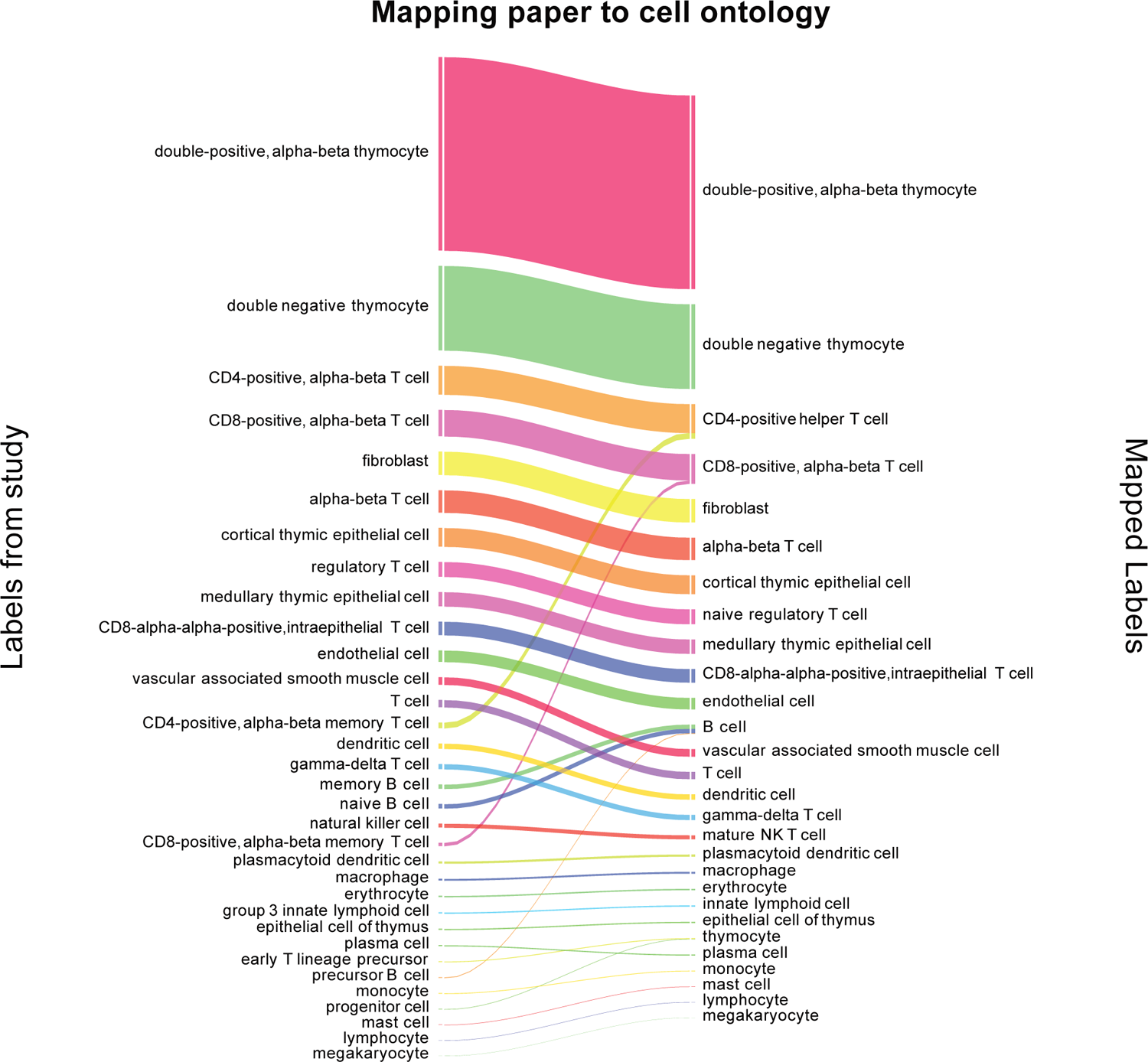
Translation of original cell type label to ontology cell type labels for thymus query data set. Cell types were relabeled based on this alluvial plot to assign them to cell ontology terms and make the granularity comparable between reference and query data set. For every cell type, the closest match in the reference data set was identified and used as the new label, if no appropriate term was found the label remained unchanged. We summed up all different types of B cells and fine sub-types of CD4 and CD8 T cells.

**Supplementary Figure 13:**
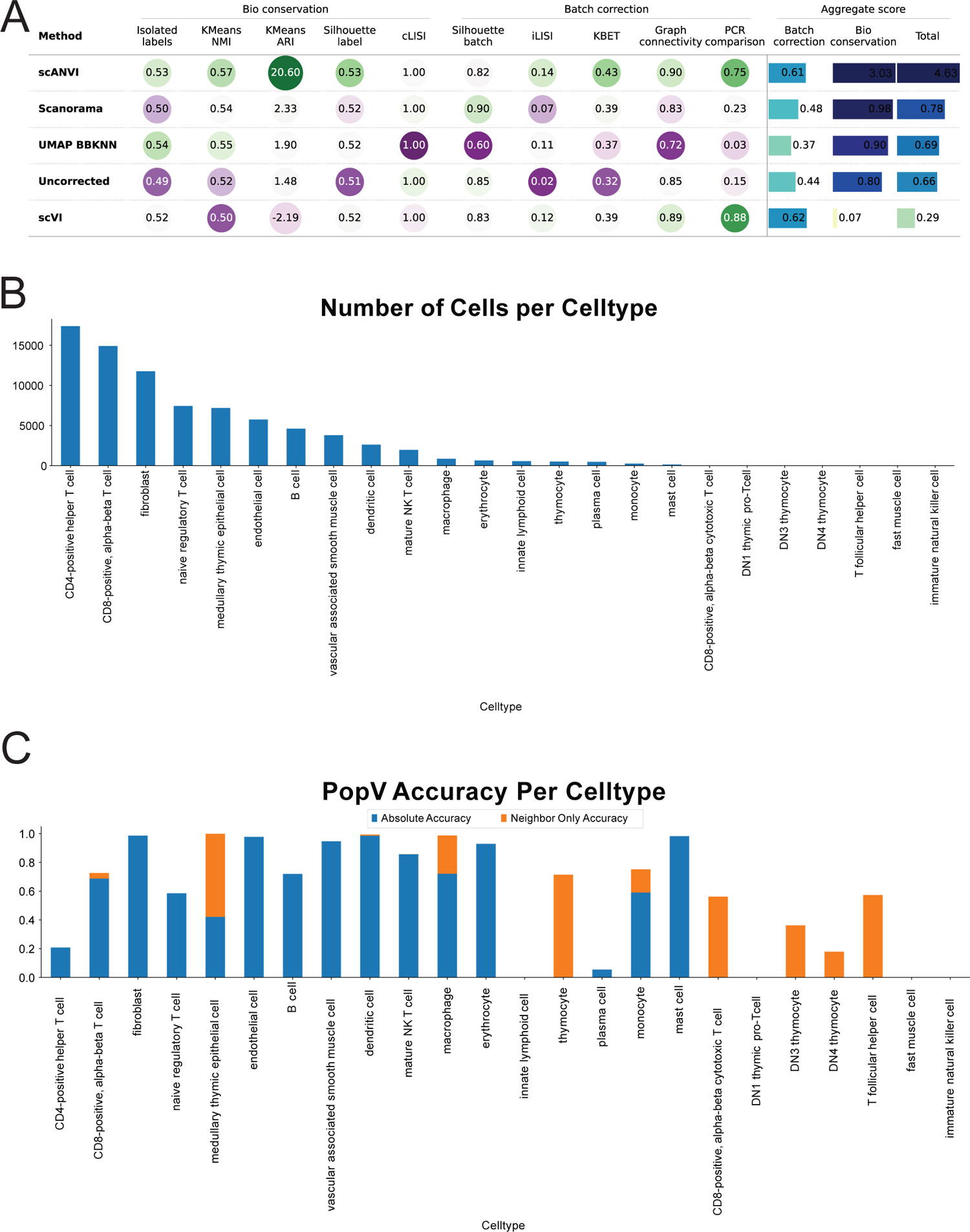
scANVI shows the highest integration of query cells and popV shows low confidence for developing T cells. **A.** scIB metrics comparing integration scores after integrating query and reference data sets showed the best integration using scANVI and improvement over uncorrected data. Labels from the original paper were used to compute cell-type dependent scores. **B, C.** Displayed is the number of each predicted cell type in query cells and the accuracy for each annotated cell type. Of note, T cells are highly abundant but not accurately predicted.

**Supplementary Figure 14:**
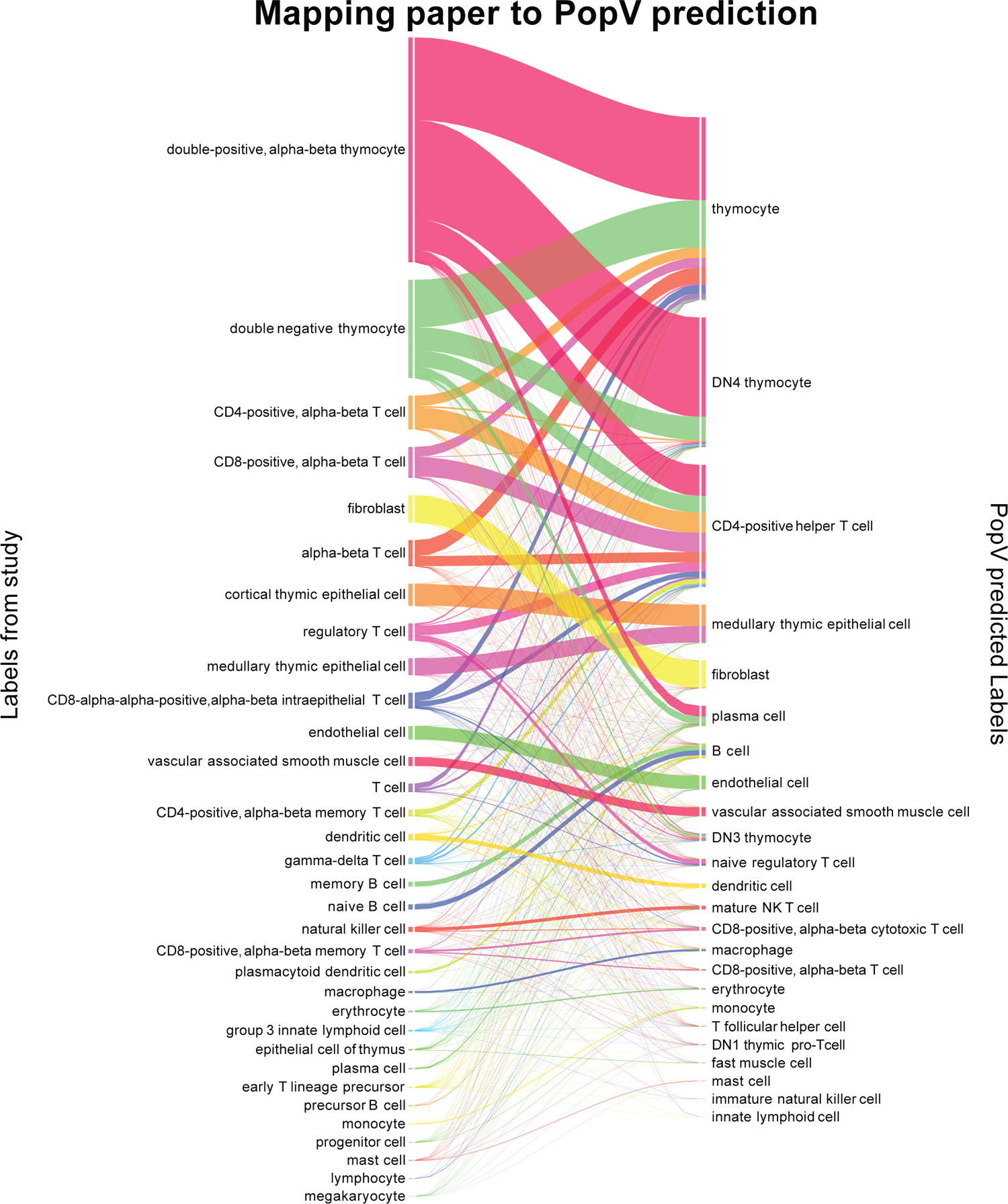
Alluvial plot of annotation in Thymus cells highlights cells with major disagreement. The major disagreement between popV prediction and original annotation in the paper is displayed. A wide variety of labels is predicted for thymocytes as well as mature T cells.

**Supplementary Figure 15:**
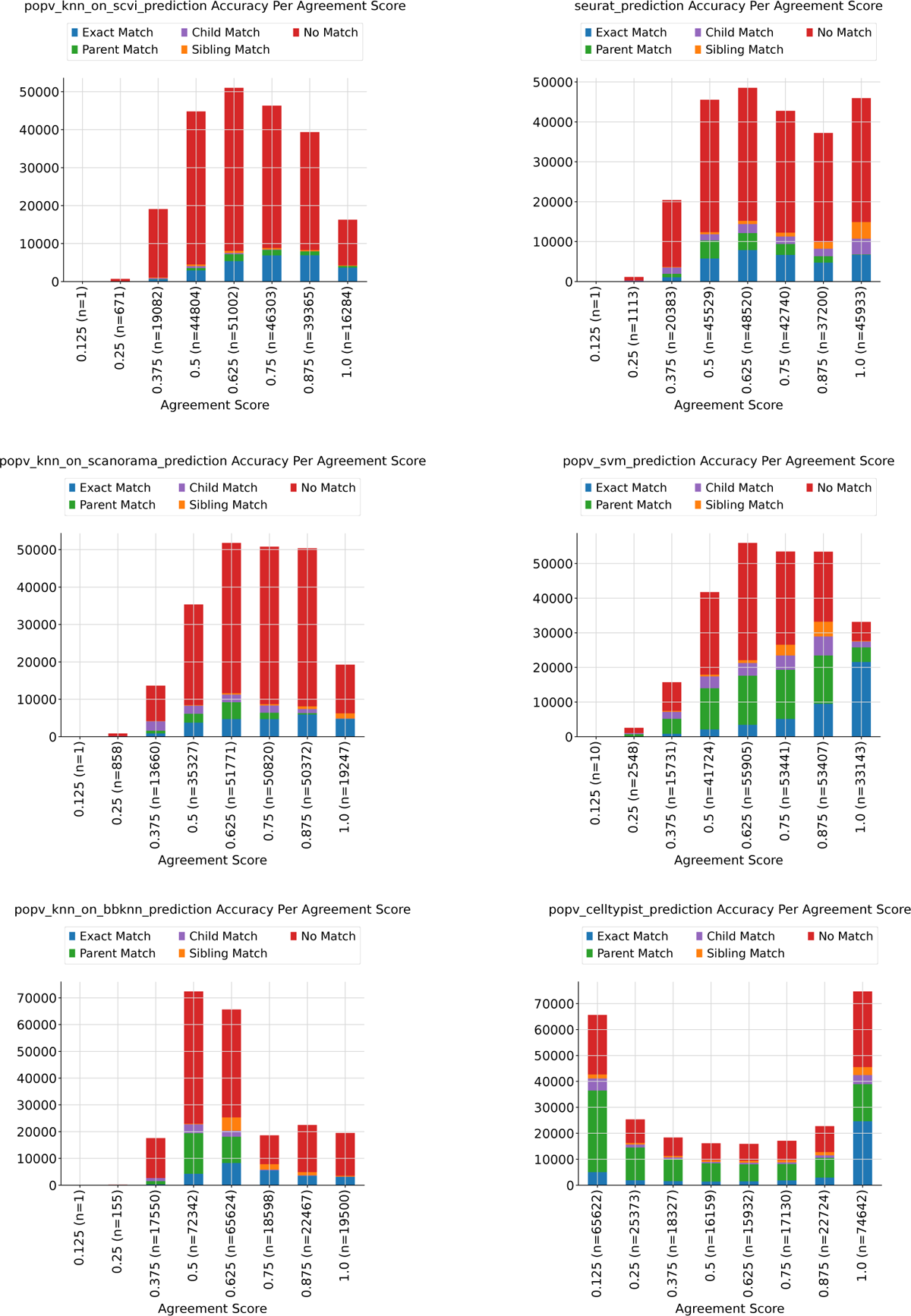
Calibration of certainty and accuracy for methods left out in 4. KNN after scVI, KNN after SCANORAMA, Seurat prediction and KNN after BBKNN integration shows no tendency to higher accuracy with higher certainty. Accuracy and certainty are almost independent of those. Celltypist shows a high proportion of low-probability as well as high-probability cells and a high number of parent matches. Probabilities and ground truth are not correlated. SVM shows a good correlation between accuracy and certainty with an 80% accuracy for high certainty prediction. This is low compared to the PopV prediction score.

**Supplementary Figure 16:**
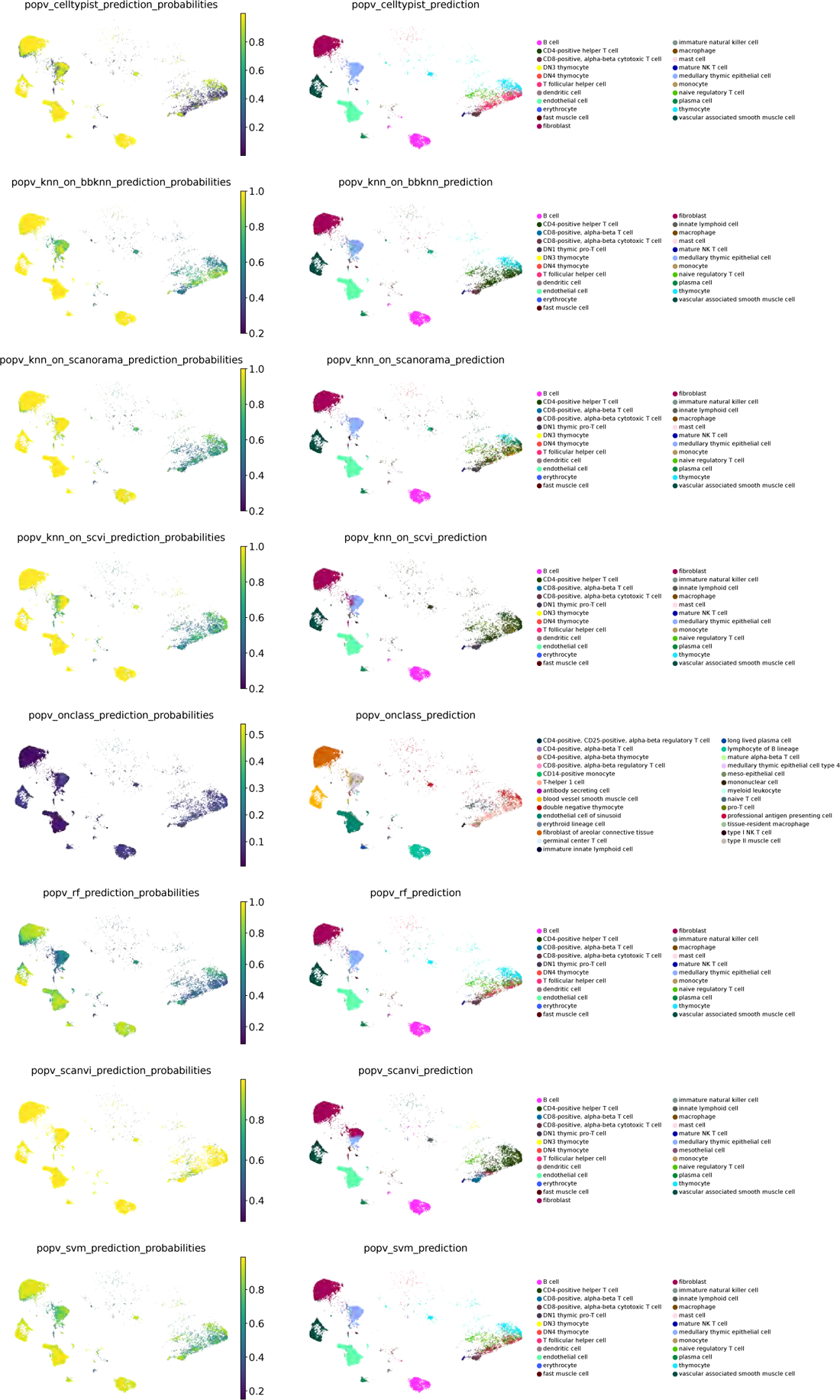
Annotation certainty and accuracy in adult thymus cells for the different algorithms. We display the internal certainty of the various algorithms compared to the respective accuracy and see again a high variability in the prediction certainty with scANVI being over-confident and RF having low certainty in its predictions. All classifiers show lower certainty for T cells compared to other cell types. Most algorithms show a lower intrinsic certainty for cortical epithelial cells (BBKNN, RF, SVM).

**Supplementary Figure 17:**
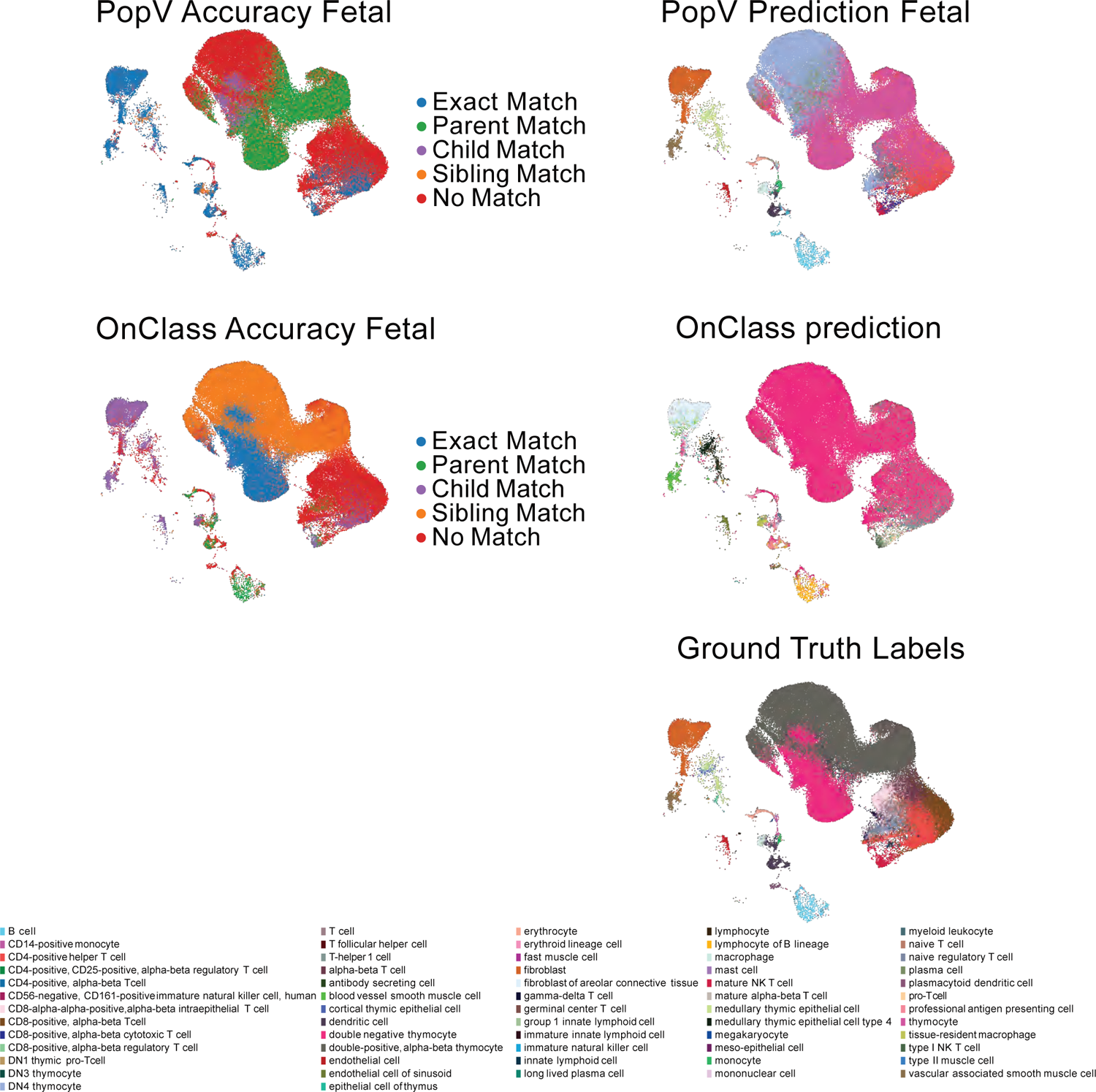
Prediction of cell types in fetal thymus using adult thymus as reference. We compare the prediction of popV with the prediction of OnClass. The reason is that OnClass can predict unseen cell types and those known query cell types are largely not seen in the reference data set. OnClass prediction shows a higher number of *Exact Match* and *Sibling Match* for thymocytes compared to PopV prediction. While PopV shows *Exact Match* for several cell types outside of the thymocytes, where OnClass predictions predicts cell-types with a child or parent relationship to the original annotation. OnClass is inaccurate for the transitional states in developing thymocytes annotating all cells as *DN thymocytes* and ignoring *DP thymocytes* as well as early mature cells. PopV predicts *DN4 thymocytes* for cells originally labeled as *DP thymocytes* as well as a large number of *thymocytes*. By construction *DN4 thymocytes* leads to *No Match* compared to the correct label *double-positive, alpha-beta thymocyte*, whereas OnClass predictions of *DN thymocytes* leads to a sibling match.

## Supplementary Tables

**Supplementary Table 1:** Removed cell-type from Tabula sapiens due to less than 10 cells per cell type.

| Tissue | Cell type |
| --- | --- |
| Eye | retina horizontal cell |
| Lymph Node | plasmacytoid dendritic cell |
| Tongue | Schwann cell |
| Blood | plasmablast |
| Uterus | B cell |
| Skin | naive B cell |
| Bone Marrow | plasmablast |
| Pancreas | pancreatic D cell |
| Fat | mast cell |
| Thymus | myeloid dendritic cell |
| Trachea | double-positive, alpha-beta thymocyte |
| Prostate | mast cell |
| Lung | myofibroblast cell |

**Supplementary Table 2:** Removed tissues from Tabula sapiens due to inconsistent cell-type annotation across donors.

| Tissue | Cell type |
| --- | --- |
| TSP14 | Liver |
| TSP2 | Trachea |
| TSP2 | Lymph Node |
| TSP2 | Spleen |
| TSP2 | Large Intestine |
| TSP2 | Small Intestine |
| TSP1 | Blood |
| TSP2 | Blood |
| TSP14 | Prostate |
| TSP2 | Thymus |

**Supplementary Table 3:** Overview of donor ID and assay in Tabula sapiens data. Part1.

| Tissue | Donor ID | Assay | Cell Count |
| --- | --- | --- | --- |
| Bladder | TSP1 | 10x 3' v3 | 11022 |
| Bladder | TSP14 | 10x 3' v3 | 3066 |
| Bladder | TSP1 | Smart-seq2 | 714 |
| Bladder | TSP2 | 10x 3' v3 | 9334 |
| Bladder | TSP2 | Smart-seq2 | 447 |
| Blood | TSP10 | 10x 3' v3 | 5004 |
| Blood | TSP14 | 10x 3' v3 | 6733 |
| Blood | TSP14 | 10x 5' v3 | 3229 |
| Blood | TSP8 | 10x 3' v3 | 1974 |
| Blood | TSP7 | 10x 3' v3 | 17213 |
| Blood | TSP7 | Smart-seq2 | 613 |
| Bone_Marrow | TSP11 | Smart-seq2 | 1031 |
| Bone_Marrow | TSP11 | 10x 3' v3 | 1456 |
| Bone_Marrow | TSP14 | 10x 3' v3 | 4457 |
| Bone_Marrow | TSP13 | Smart-seq2 | 536 |
| Bone_Marrow | TSP14 | 10x 5' v3 | 1396 |
| Bone_Marrow | TSP2 | 10x 3' v3 | 2698 |
| Bone_Marrow | TSP2 | Smart-seq2 | 719 |
| Eye | TSP15 | 10x 3' v3 | 5599 |
| Eye | TSP3 | 10x 3' v3 | 2163 |
| Eye | TSP3 | Smart-seq2 | 279 |
| Eye | TSP5 | 10x 3' v3 | 2453 |
| Eye | TSP5 | Smart-seq2 | 132 |
| Fat | TSP10 | Smart-seq2 | 648 |
| Fat | TSP10 | 10x 3' v3 | 11941 |
| Fat | TSP14 | 10x 3' v3 | 7670 |
| Heart | TSP12 | Smart-seq2 | 277 |
| Heart | TSP12 | 10x 3' v3 | 11228 |
| Kidney | TSP2 | 10x 3' v3 | 9271 |
| Kidney | TSP2 | Smart-seq2 | 370 |
| Large_Intestine | TSP14 | 10x 3' v3 | 11439 |
| Liver | TSP6 | 10x 3' v3 | 2647 |
| Liver | TSP6 | Smart-seq2 | 213 |
| Lung | TSP14 | 10x 3' v3 | 4350 |
| Lung | TSP1 | 10x 3' v3 | 12507 |
| Lung | TSP1 | Smart-seq2 | 689 |
| Lung | TSP2 | 10x 3' v3 | 17225 |
| Lung | TSP2 | Smart-seq2 | 901 |
| Lymph_Node | TSP14 | 10x 3' v3 | 16383 |
| Lymph_Node | TSP14 | 10x 5' v3 | 14044 |
| Lymph_Node | TSP7 | Smart-seq2 | 1162 |
| Lymph_Node | TSP7 | 10x 3' v3 | 11913 |
| Mammary | TSP4 | 10x 3' v3 | 10795 |
| Mammary | TSP4 | Smart-seq2 | 580 |
| Muscle | TSP14 | 10x 3' v3 | 8394 |
| Muscle | TSP1 | 10x 3' v3 | 2512 |
| Muscle | TSP1 | Smart-seq2 | 1248 |
| Muscle | TSP2 | 10x 3' v3 | 14797 |
| Muscle | TSP2 | Smart-seq2 | 1550 |
| Muscle | TSP4 | Smart-seq2 | 2245 |
| Pancreas | TSP9 | 10x 3' v3 | 6616 |
| Pancreas | TSP9 | Smart-seq2 | 83 |
| Pancreas | TSP1 | 10x 3' v3 | 5876 |
| Pancreas | TSP1 | Smart-seq2 | 913 |
| Prostate | TSP8 | Smart-seq2 | 633 |
| Prostate | TSP8 | 10x 3' v3 | 12460 |

**Supplementary Table 4:** Overview of donor ID and assay in Tabula sapiens data. Part2.

| Tissue | Donor ID | Assay | Cell Count |
| --- | --- | --- | --- |
| Salivary_Gland | TSP14 | 10x 3' v3 | 17762 |
| Salivary_Gland | TSP7 | Smart-seq2 | 832 |
| Salivary_Gland | TSP7 | 10x 3' v3 | 8605 |
| Skin | TSP10 | Smart-seq2 | 839 |
| Skin | TSP10 | 10x 3' v3 | 5008 |
| Skin | TSP14 | 10x 3' v3 | 3551 |
| Small_Intestine | TSP14 | 10x 3' v3 | 10458 |
| Spleen | TSP14 | 10x 3' v3 | 10935 |
| Spleen | TSP14 | 10x 5' v3 | 8029 |
| Spleen | TSP7 | Smart-seq2 | 1167 |
| Spleen | TSP7 | 10x 3' v3 | 6194 |
| Thymus | TSP14 | 10x 3' v3 | 8729 |
| Thymus | TSP14 | 10x 5' v3 | 12846 |
| Tongue | TSP14 | 10x 3' v3 | 5407 |
| Tongue | TSP7 | Smart-seq2 | 1205 |
| Tongue | TSP4 | Smart-seq2 | 186 |
| Tongue | TSP7 | 10x 3' v3 | 8212 |
| Trachea | TSP6 | 10x 3' v3 | 4757 |
| Trachea | TSP6 | Smart-seq2 | 355 |
| Uterus | TSP4 | Smart-seq2 | 286 |
| Uterus | TSP4 | 10x 3' v3 | 6828 |
| Vasculature | TSP14 | 10x 3' v3 | 7389 |
| Vasculature | TSP2 | 10x 3' v3 | 7801 |
| Vasculature | TSP2 | Smart-seq2 | 847 |

**Supplementary Table 5:** Overview of donor ID and assay in Lung Cell Atlas.

| Donor | Sample | Assay | Tissue | Cell Count |
| --- | --- | --- | --- | --- |
| 1 | blood 1 | 10x 3' v2 | blood | 2220 |
| 3 | blood 3 | 10x 3' v2 | blood | 2449 |
| 1 | blood 1 | Smart-seq2 | blood | 752 |
| 3 | proximal 3 | 10x 3' v2 | lung | 7773 |
| 3 | medial 3 | Smart-seq2 | lung | 559 |
| 3 | distal 3 | 10x 3' v2 | lung | 16903 |
| 3 | distal 3 | Smart-seq2 | lung | 1190 |
| 2 | distal 2 | 10x 3' v2 | lung | 18542 |
| 1 | distal 1b | Smart-seq2 | lung | 1642 |
| 1 | distal 1a | Smart-seq2 | lung | 1593 |
| 1 | distal 1a | 10x 3' v2 | lung | 7524 |
| 2 | medial 2 | Smart-seq2 | lung | 1291 |
| 2 | medial 2 | 10x 3' v2 | lung | 10251 |
| 2 | distal 2 | Smart-seq2 | lung | 1982 |
| 3 | proximal 3 | Smart-seq2 | lung | 400 |

**Supplementary Table 6:**
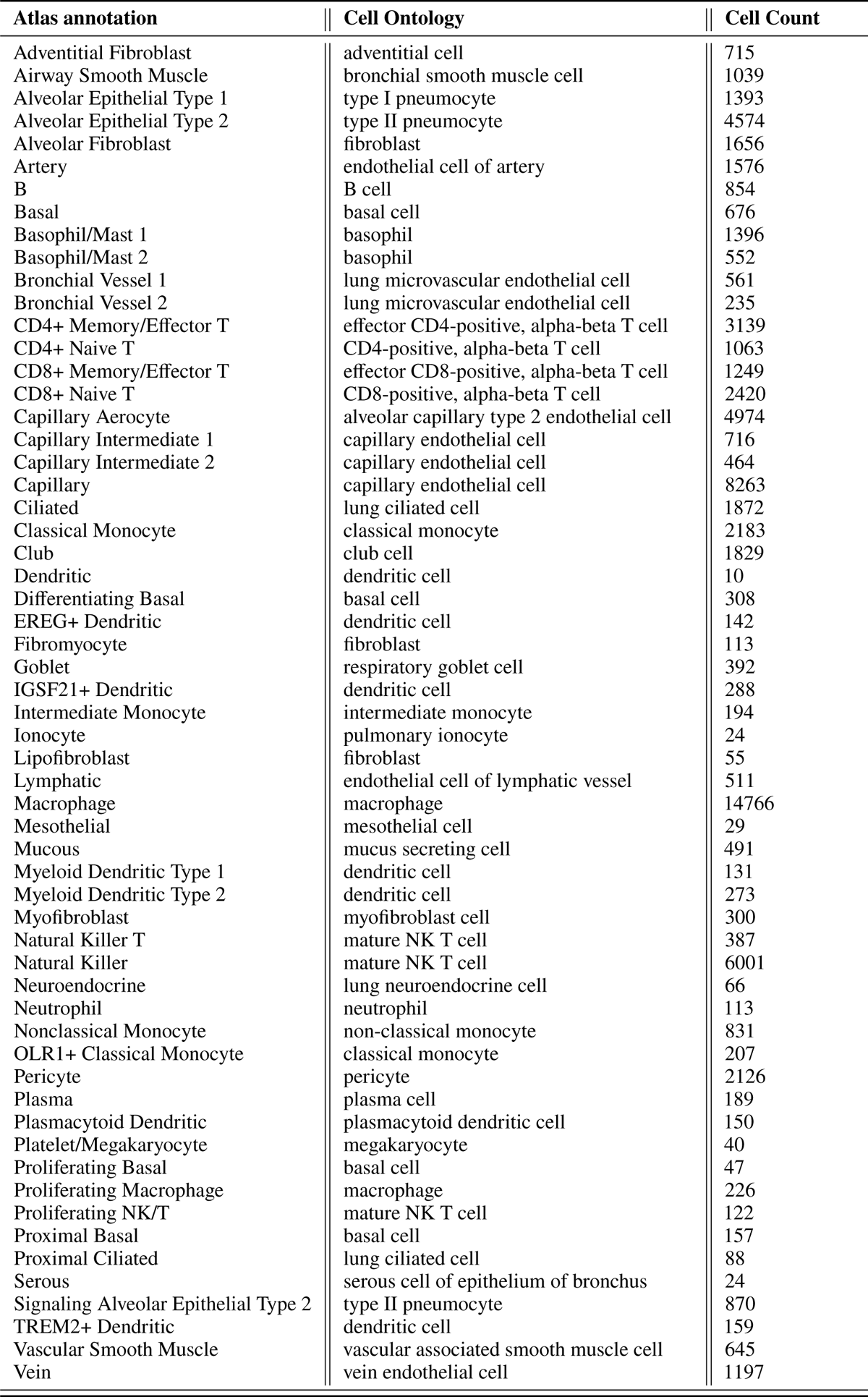
Mapping of lung cell atlas annotations to annotations for cell ontology and accuracy computation.

**Supplementary Table 7:**
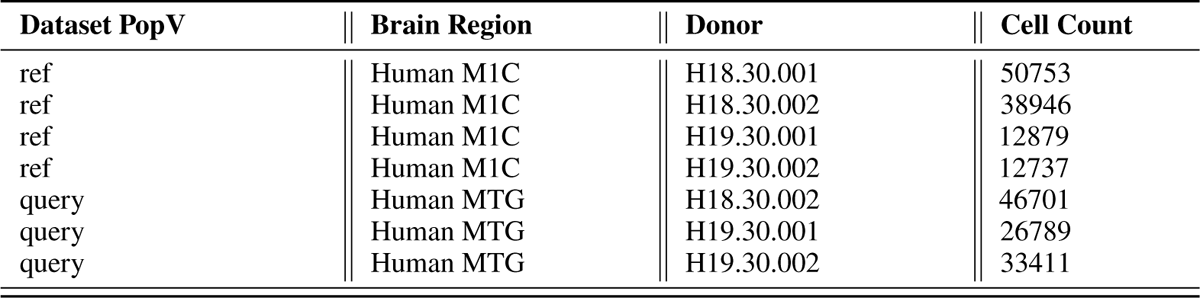
Overview of donor ID and assay in both brain data sets.

| Dataset PopV | Brain Region | Donor | Cell Count |
| --- | --- | --- | --- |
| ref | Human M1C | H18.30.001 | 50753 |
| ref | Human M1C | H18.30.002 | 38946 |
| ref | Human M1C | H19.30.001 | 12879 |
| ref | Human M1C | H19.30.002 | 12737 |
| query | Human MTG | H18.30.002 | 46701 |
| query | Human MTG | H19.30.001 | 26789 |
| query | Human MTG | H19.30.002 | 33411 |

**Supplementary Table 8:** Overview of donor ID and assay in Thymus data set.

| Development Stage | Donor | Assay | Cell Count |
| --- | --- | --- | --- |
| human early adulthood stage | A43 | 10x 5' v1 | 22804 |
| young adult stage | A16 | 10x 5' v1 | 11491 |
| young adult stage | A16 | 10x 3' v2 | 11180 |
| organogenesis stage | C40 | 10x 3' v3 | 14021 |
| organogenesis stage | C41 | 10x 3' v3 | 11409 |
| 9th week post-fertilization human stage | C34 | 10x 5' v1 | 390 |
| 9th week post-fertilization human stage | F22 | 10x 3' v2 | 3137 |
| 10th week post-fertilization human stage | F74 | 10x 5' v1 | 7410 |
| 11th week post-fertilization human stage | F64 | 10x 5' v1 | 7739 |
| 11th week post-fertilization human stage | F23 | 10x 3' v2 | 5774 |
| 12th week post-fertilization human stage | F45 | 10x 3' v2 | 10380 |
| 12th week post-fertilization human stage | F45 | 10x 5' v1 | 9630 |
| 12th week post-fertilization human stage | F67 | 10x 5' v1 | 13579 |
| 13th week post-fertilization human stage | F38 | 10x 5' v1 | 3047 |
| 13th week post-fertilization human stage | F38 | 10x 3' v2 | 5987 |
| 14th week post-fertilization human stage | F30 | 10x 3' v2 | 7535 |
| 14th week post-fertilization human stage | F30 | 10x 5' v1 | 5199 |
| 16th week post-fertilization human stage | F41 | 10x 5' v1 | 3363 |
| 16th week post-fertilization human stage | F41 | 10x 3' v2 | 6238 |
| 16th week post-fertilization human stage | F21 | 10x 3' v2 | 8481 |
| 17th week post-fertilization human stage | F83 | 10x 5' v1 | 3678 |
| 17th week post-fertilization human stage | F29 | 10x 3' v2 | 6146 |
| 17th week post-fertilization human stage | F29 | 10x 5' v1 | 5254 |
| infant stage | P3 | 10x 3' v2 | 12288 |
| infant stage | P1 | 10x 3' v2 | 10811 |
| adolescent stage | P2 | 10x 3' v2 | 12423 |
| 2-year-old human stage | T06 | 10x 5' v1 | 19423 |
| 3-month-old human stage | T07 | 10x 5' v1 | 12295 |
| 10-month-old human stage | T03 | 10x 5' v1 | 4789 |

**Supplementary Table 9:** Mapping of thymus annotations to annotations for cell ontology and accuracy computation.

| Atlas annotation | Cell Ontology | Cell Count |
| --- | --- | --- |
| CD8-positive, alpha-beta T cell | CD8-positive, alpha-beta T cell | 13257 |
| CD4-positive, alpha-beta T cell | CD4-positive helper T cell | 14506 |
| double-positive, alpha-beta thymocyte | double-positive, alpha-beta thymocyte | 97183 |
| CD8-alpha-alpha-positive, alpha-beta intraepithelial T cell | CD8-alpha-alpha-positive, alpha-beta intraepithelial T cell | 6783 |
| regulatory T cell | naive regulatory T cell | 7444 |
| alpha-beta T cell | alpha-beta T cell | 11235 |
| T cell | T cell | 3614 |
| double negative thymocyte | double negative thymocyte | 42474 |
| monocyte | monocyte | 265 |
| naive B cell | B cell | 2152 |
| dendritic cell | dendritic cell | 2625 |
| CD4-positive, alpha-beta memory T cell | CD4-positive helper T cell | 2879 |
| gamma-delta T cell | gamma-delta T cell | 2582 |
| plasmacytoid dendritic cell | plasmacytoid dendritic cell | 1048 |
| natural killer cell | mature NK T cell | 1964 |
| macrophage | macrophage | 863 |
| memory B cell | B cell | 2161 |
| precursor B cell | B cell | 290 |
| progenitor cell | thymocyte | 185 |
| fibroblast | fibroblast | 11749 |
| early T lineage precursor | thymocyte | 316 |
| mast cell | mast cell | 148 |
| group 3 innate lymphoid cell | innate lymphoid cell | 561 |
| erythrocyte | erythrocyte | 644 |
| vascular associated smooth muscle cell | vascular associated smooth muscle cell | 3788 |
| medullary thymic epithelial cell | medullary thymic epithelial cell | 7193 |
| epithelial cell of thymus | epithelial cell of thymus | 554 |
| endothelial cell | endothelial cell | 5753 |
| cortical thymic epithelial cell | cortical thymic epithelial cell | 9411 |
| megakaryocyte | megakaryocyte | 36 |
| lymphocyte | lymphocyte | 115 |
| plasma cell | plasma cell | 479 |
| CD8-positive, alpha-beta memory T cell | CD8-positive, alpha-beta T cell | 1644 |

## Data availability

All pre-trained model are located at https://doi.org/10.5281/zenodo.7580707. All pre-processed data objects used to pretrain the Tabula sapiens reference models are deposited at https://doi.org/10.5281/zenodo.7587774. All other data is freely accessible through CELLx-GENE.

## Code availability

The code to reproduce the experiments of this manuscript will be made available upon publication at https://github.com/YosefLab/popv-reproducibility. The PopV package can be found on GitHub at https://github.com/czbiohub/PopV,

## Notes

https://zenodo.org/record/7580707#.ZBpRUS-B19g

### Competing Interest Statement

Nir Yosef is an advisor and/or has equity in Cellarity, Celsius Therapeutics, and Rheos Medicine.

